# Limited haplotype diversity underlies polygenic trait architecture across 70 years of wheat breeding

**DOI:** 10.1101/2020.09.15.296533

**Authors:** Michael F. Scott, Nick Fradgley, Alison R. Bentley, Thomas Brabbs, Fiona Corke, Keith A. Gardner, Richard Horsnell, Phil Howell, Olufunmilayo Ladejobi, Ian J. Mackay, Richard Mott, James Cockram

## Abstract

**Background:** Breeding has helped improve bread wheat yield significantly over the last century. Understanding the potential for future crop improvement depends on relating segregating genetic variation to agronomic traits.

**Results:** We bred NIAB Diverse MAGIC population, comprising over 500 recombinant inbred lines, descended from sixteen bread wheat varieties released between 1935-2004. We sequenced the founders’ exomes and promotors by capture. Despite being highly representative of North-West European wheat and capturing 73% of global polymorphism, we found 89% of genes contained no more than three haplotypes. We sequenced each line with 0.3x coverage whole-genome sequencing, and imputed 1.1M high-quality SNPs that were over 99% concordant with array genotypes. Imputation accuracy remained high at coverage as low as 0.076x, with or without the use of founder genomes as reference panels. We created a genotype-phenotype map for 47 traits over two years. We found 136 genome-wide significant associations, concentrated at 42 genetic loci with large and often pleiotropic effects. Outside of these loci most traits are polygenic, as revealed by multi-locus shrinkage modelling.

**Conclusions:** Historically, wheat breeding has reshuffled a limited palette of haplotypes; continued improvement will require selection at dozens of loci of diminishing effect, as most of the major loci we mapped are known. Breeding to optimise one trait generates correlated trait changes, exemplified by the negative trade-off between yield and protein content, unless selection and recombination can break critical unfavourable trait-trait associations. Finally, low coverage whole genome sequencing of bread wheat populations is an economical and accurate genotyping strategy.

## Introduction

Bread wheat (*Triticum aestivum* L.) production is a critical component of worldwide food security. Demand for wheat is predicted to increase by 60% between 2014 and 2050[1], by which time the human population will have reached 9 billion. Breeding will be a key component of meeting this demand sustainably[2]. Over the past century, genetic gains have been responsible for between one third and two thirds of yield improvements in European wheats, amounting to a 12-120kg increase in yield (~1%) per hectare per year[3–6].

Genomic data is expected to accelerate the rate of genetic gain in wheat[7,8]. Surveys of global standing genetic variation include, for example, whole genome resequencing (WGS) of 93 accessions[9], exome capture for 870 accessions[10], genotyping by sequencing (~16k markers) for ~17k breeding programme lines[11], and genotyping array data for collections of 804[12] and 4,500[13] accessions (~15k and ~113k markers, respectively). Bread wheat’s large genome size (17Gb) inflates the cost of collecting sequencing data and its hexaploidy reduces the accuracy and cross-population consistency of genotyping array data[12]. The potential for genotyping by low-coverage WGS in polyploid wheat has yet to be established.

To aid genetic gain through breeding, it is crucial to link genetic data with phenotypic information and thereby reveal genotype-phenotype associations[11,14]. Previous genotypic/phenotypic datasets include five traits measured in two years for 870 global accessions with exome capture data[10], and 12 traits measured in two years, six locations, and three cropping intensities for 191 German varieties with genotyping array data (~9k markers)[15]. Genotype-trait and trait-trait associations may be confounded by population structure or hidden by low allele frequencies in studies of existing varieties or breeding lines. These problems can be controlled in experimental populations produced by crossing. However, mapping resolution and overall genetic diversity are typically low in experimental populations. Multiparent Advanced Generation Intercross (MAGIC) populations are designed to address these issues by accumulating recombination events through generations of intercrossing and capturing diversity across multiple founders[16–18].

In this study we undertook a systematic approach to these challenges. We bred a new multi-parental population, the ‘NIAB DIVERSE MAGIC’ population (hereafter ‘NDM’) through hundreds of structured inter-crosses between sixteen diverse founders. Our multi-funnel crossing design creates a greater number and more uniform genome-wide distribution of recombinant haplotypes than alternative multiparent populations[19] and the relatively large number of diverse founders samples more genetic variation. We sampled founders released between 1935-2004, aiming to determine the genetic basis for historical changes in agronomic traits and the potential for future improvement from within the existing pool of variants. We used a cost-effective genotyping strategy by low-coverage WGS, accurately imputing over 1M SNPs in over 500 recombinant inbred lines. We measured 47 phenotypes in the population, of which 25 were assessed across two growing seasons. The power of NDM comes from the combination of carefully designed germplasm and dense genotypic and phenotypic information, all of which we make publicly available.

We address the following questions. First, what genetic variation exists among the exomes of the NDM founders, how does it reflect global wheat diversity, and specifically how many distinct haplotypes typically segregate at each locus? Second, how does this variation underlie agronomic traits, as revealed through genetic mapping and genomic prediction? And third, what do these models imply about the future potential for phenotypic change and to what extent should we expect selection to cause correlated trait changes due to the sharing of causal genetic variants between traits.

## Results

### NIAB DIVERSE MAGIC Founders

The 16 founders were selected from a panel of 94 historical varieties released in the UK over a ~70 year period (and originating from the UK, France, Denmark, Sweden and the Netherlands, Supplementary Table 1) using 546 Diversity Array Technology (DArT) and 61 Single Sequence Repeat (SSR) markers[20]. We sequenced 15 founders after enrichment for (a) genic regions and (b) putative promoters using a capture probe-set[21] at average coverage of 22.94x of the targets (Supplementary Table 1). The remaining founder, Holdfast, was sequenced by WGS, but to ensure consistency across founders, we restricted our attention to the capture targets, for which coverage in Holdfast was 15.8x. We sequenced using Illumina 150bp paired-end reads whose combined length often included sequence differences between homeologous loci on the A, B and D subgenomes of hexaploid wheat, thereby resolving otherwise ambiguous alignments. Furthermore, we only used high quality alignments (mapQ>30) for coverage calculations and variant calling, and subsequently excluded variant sites with missing or heterozygous calls in any founder (e.g. from homeologous variation and misalignment). After quality control, we called 1.13M high-quality single nucleotide polymorphisms (SNPs) across the 110,790 promoter-gene pairs targeted by the capture probes (557Mb in total), summarised in Supplementary Figure 1. Only 97,727 SNPs (8.7%) were on the D subgenome and almost half (17,289/35,021, 49.4%) of the promoter-gene pairs on the D subgenome had no SNPs passing quality control, compared to 26.6% (9,656/36,302) and 21.7% (8,012/36,738) on the A and B subgenomes, respectively. A comparative lack of diversity is expected on the D subgenome as it was acquired in the most recent allo-polyploidisation event[22].

We placed the 16 founders in the context of global wheat diversity by analysing 113,457 genotyping array sites that vary among 4,506 diverse global wheat accessions[13], of which 50,335 sites were callable across all founders. We classified global wheats into nested subsets representing the UK only (n=154), North-West (NW) Europe (n=1,343), Europe (n=2,331), and Global (n=4,506), to understand how allele frequencies across subsets relate to our founders (Figure 1). Most Global common variants are polymorphic in the founders whereas rare alleles are more likely to be fixed in the founders, particularly those scarce in NW Europe and the UK. For example, 79.7% of those SNPs polymorphic within the UK subset (which includes landraces) also segregate among the founders, falling to 73.4% Global sites across all 4,506 accessions. We next asked whether we could have selected 16 founders that more comprehensively sampled the variation space. We simulated selections from the same nested subsets and compared the distribution of the fraction of segregating sites with that in the actual NDM founders, and found the latter capture more diversity than an average selection of UK wheats, about average diversity for NW European wheats, but less than average for wider European and Global sets (Figure 1). As the Global dataset is highly diverse, with modern varieties (released 1960-2009, n=2,294), landraces (1800-1959, n=965), and uncategorised/landrace germplasm (n=1,247), we conclude that NDM is representative of NW European wheat germplasm.

**Figure 1.**
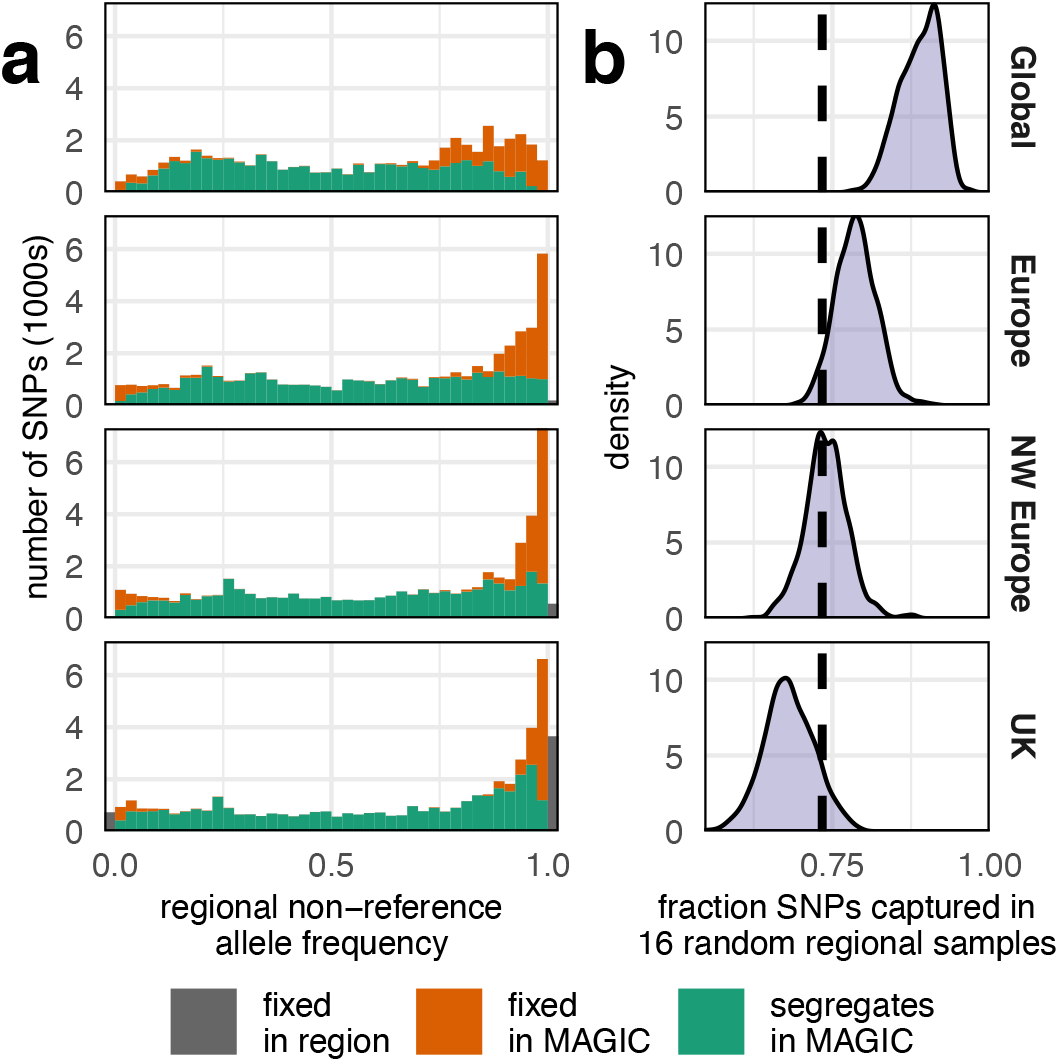
The NDM population is representative of NW European wheat. (a) SNPs segregating (green) or fixed (orange) in NDM at 50,335 sites in 4,506 global wheats, grouped into ‘Global’, ‘European’, ‘North-West European’ and ‘UK’ nested subpopulations and binned by the allele frequency in these subpopulations. (b) The fraction of sites that are polymorphic in 16 randomly chosen wheats from each subpopulation based on 1000 random replications. The dashed vertical black line at x=0.734 is the fraction of SNPs segregating among NDM founders.

We next estimated the haplotypic diversity in the founders at the 1.13M sites. First, we clustered the founders by their haplotypic similarity at the 73,982/110,790 (66.7%) promoter-gene loci with at least two haplotypes. Assuming that founders carry the same haplotype when their genotypic similarity exceeds 95%, 38,535 loci (52% of loci with variants) had only two haplotypes, 61,438 loci (83%) had at most three haplotypes, and 70,602 loci (95%) had four haplotypes at most (Figure 2b). Second, we estimated haplotype diversity by a dynamic programming algorithm that adjusted locus/block boundaries (Figure 2c, Supplementary Figure 2) to minimise the number of distinct haplotypes within a locus, while balancing transitions between calling identical versus non-identical haplotypes. Over a wide range of parameters, the average number of haplotypes present at any locus rarely exceeded two (Supplementary Figure 2: 81.2% of 1.13M sites inferred to have just two haplotypes). This analysis found slightly fewer haplotypes than the gene-based analysis because it can infer one haplotype (4.1% of sites) when nearby variation is inconsistent, and split genes with high haplotypic diversity into multiple blocks.

**Figure 2.**
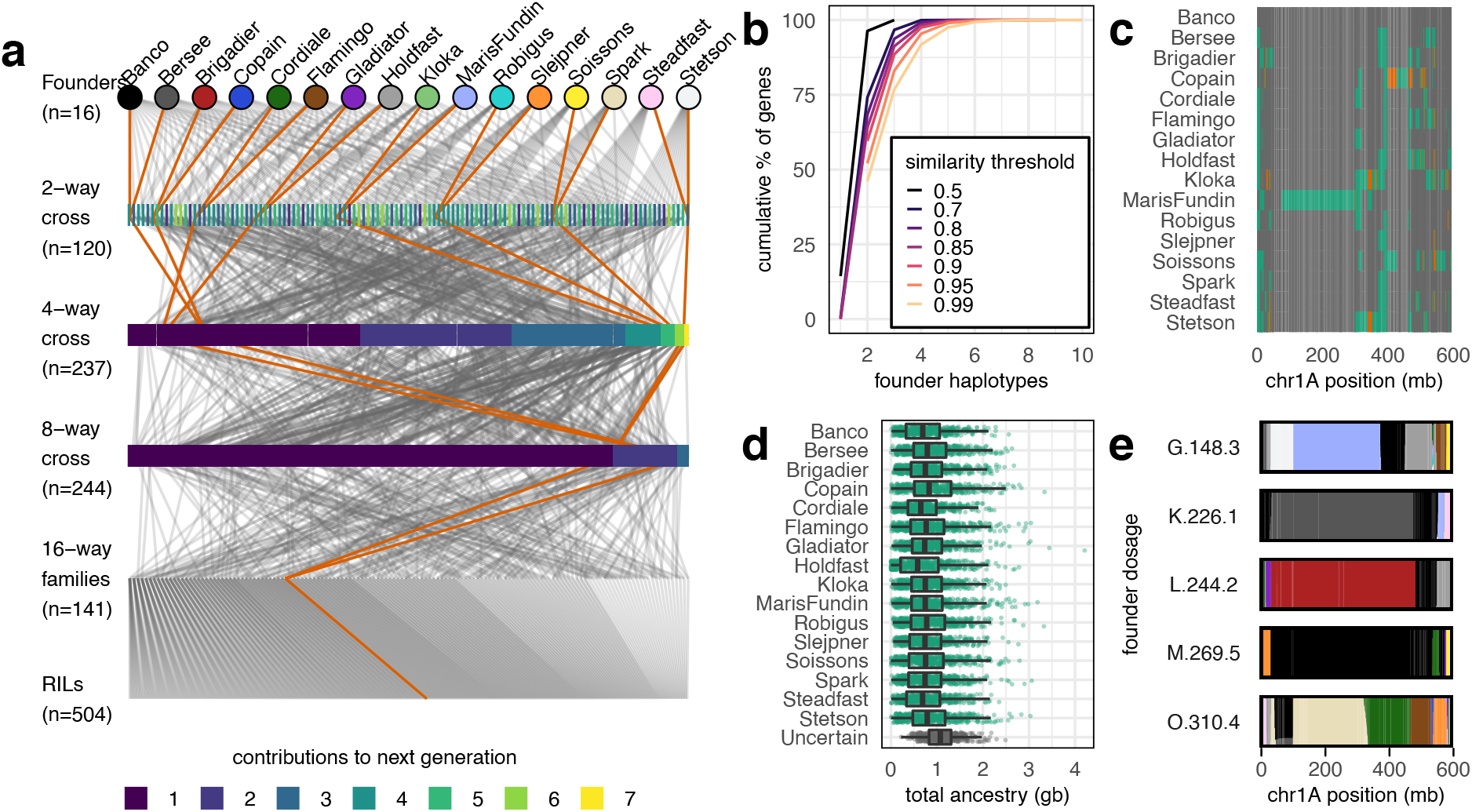
NDM population design and haplotypic diversity. (a) Pedigree showing the construction of 504 Recombinant Inbred Lines (RILs). One exemplar pedigree is highlighted to show how all 16 founders are intercrossed into each RIL. (b) Founder haplotype groups at 73,982 promoter-gene loci with SNP variation, where founders with the same haplotype have genotypic similarity fractions that exceed the corresponding threshold. (c) Pairwise similarity/dissimilarity between founders on chromosome 1A, determined using a dynamic programming algorithm to infer founder similarity and breakpoint position. Founders that are inferred to have similar haplotypes for each region are the same colour. (d) The total length of genomic blocks in NDM lines inferred to come from each founder; uncertain ancestry blocks have a maximum founder dosage of <90%. (e) Inferred founder dosage and ancestry mosaics across chromosome 1A for five example RILs, with founders coloured as in (a).

For comparison, the 19 natural accessions that founded the *Arabidopsis thaliana* MAGIC[23] display much greater haplotypic polymorphism[24]. In *A. thaliana*, genic haplotypes were determined at the level of protein sequence similarity (>95% similarity within haplotypes)[24]. On average there were 4.8 protein haplotypes per gene and 7,263/13,919 (52.2%) of genes with two (n=4,825) or three (n=2,438) haplotypes (excluding genes with no variation). Our estimates for the NDM founders are 2.7 haplotypes per gene and 83% of variable genes having at most three haplotypes. Protein-level differences are lower than DNA level differences making this comparison conservative, and thus the true difference is even greater.

### The NIAB DIVERSE MAGIC Population

We developed a total of 596 Recombinant Inbred Lines (RILs), each descended from all 16 founders via a crossing funnel (Figure 2a). After 6 generations of inbreeding, all 596 RILs were initially genotyped using the Axiom 35k wheat breeders’ SNP genotyping array[12]. We called SNPs at 20,688 sites, of which 5,747 overlapped with the 1.13M SNP calls made in the founders. These overlapping sites suggested that only 59.8% of genotyping array probes could have been unambiguously placed using BLASTn[25], underlining the difficulty of using short probes in polyploids (Supplementary Table 2). We used the overlapping sites as a truth genotype set to find sample misidentifications and estimate the accuracy of sequence-based genotyping in the RILs.

We excluded 46 RILs excessively similar (>92%) to other RILs, indicating possible errors during population development. We sequenced the remaining 550 RILs after 7 generations of inbreeding by low coverage WGS (mean 0.304X) and called variants at the 1.13M founder SNP sites using sequence alignments. A further 46 RILs were excluded as their genotypic concordance with the initial 35k array data was below 95%, leaving 504 RILs in 141 families (RILs in the same ‘family’ are derived from the same 16-way cross), from which we based our main analyses.

We imputed RIL genotypes using STITCH[26] by inferring the founder haplotype carried by each line at each location. Figure 2c shows the haplotypic similarity among founders on chromosome 1A, indicating that a small number of haplotypes have been heavily recombined during the 69 years of breeding history that separates the founders. Most recombination is located towards the distal ends of the chromosomes, as expected[27]. Only limited further recombination occurs during MAGIC population construction and the haplotype blocks inherited from each founder are relatively long (Supplementary Figure 2) and therefore distinguishable from one another. Thus, it was necessary to assume 16 unique haplotypes were segregating to obtain the highest imputation accuracy (Supplementary Figure 2). Founder haplotypes could be confidently assigned (i.e. with >90% dosage from a single founder) at over 92.2% of sites (Figure 2d). These haplotype assignments implied that an average of 4.8-13.7 recombination events occurred per RIL per chromosome (mean 8.7 sd 2), giving an average of 183 (sd 36.3) recombination events per RIL in total. Consistent with estimated genetic map lengths of 35-37.4M[12,28], 4.9-5.2 recombination events were observed per Morgan, in line with the predicted ~5-fold increase in 16-parent MAGIC populations compared to two-way crosses[29]. Example founder haplotype mosaics across chromosome 1A are shown in Figure 2e.

The fraction of sites called directly (i.e. without imputation) for 501 RILs varied between 20.9-42.7% (mean 27.8% sd 3.4%), as expected for 0.3x-coverage sequence data. A further three RILs were sequenced to higher depth (2.7x, 4.0x, and 4.3x) and had call rates of 79.9%, 90.0%, and 93.0%, respectively (Figure 3a). After imputation, 94.2% of the 1.13M SNPs (i.e. 1.07M) were called across all 504 RILs and the effective call rate of imputed sites was 99.6%, with 5.8% of the SNP sites inaccessible or removed by quality control: 0.93% of sites are on the “Un” chromosome in the wheat reference (excluded from imputation), 1.36% were removed by imputation QC (info score <0.4) and 3.52% had imputed minor allele frequencies below 2.5% and/or missingness above 90%. Figure 3b shows that the concordance between array and imputed genotypes (AI) and between array and directly called genotypes (AD) are strongly correlated, suggesting that instances of poorer concordance are unlikely to be caused by imputation. Overall, imputation marginally improved accuracy versus direct calls (mean AI 99.1% versus mean AD 99.0%) but increased the call rate three-fold. Downsampling read coverage showed the founder haplotype space and recombination mosaics could be accurately inferred from coverage as low as 0.076x per sample (Figure 3c); above this level imputation accuracy was independent of whether founder haplotypes were included as a reference panel (mean AI 98.7%) or ignored (mean AI 98.5%).

**Figure 3.**
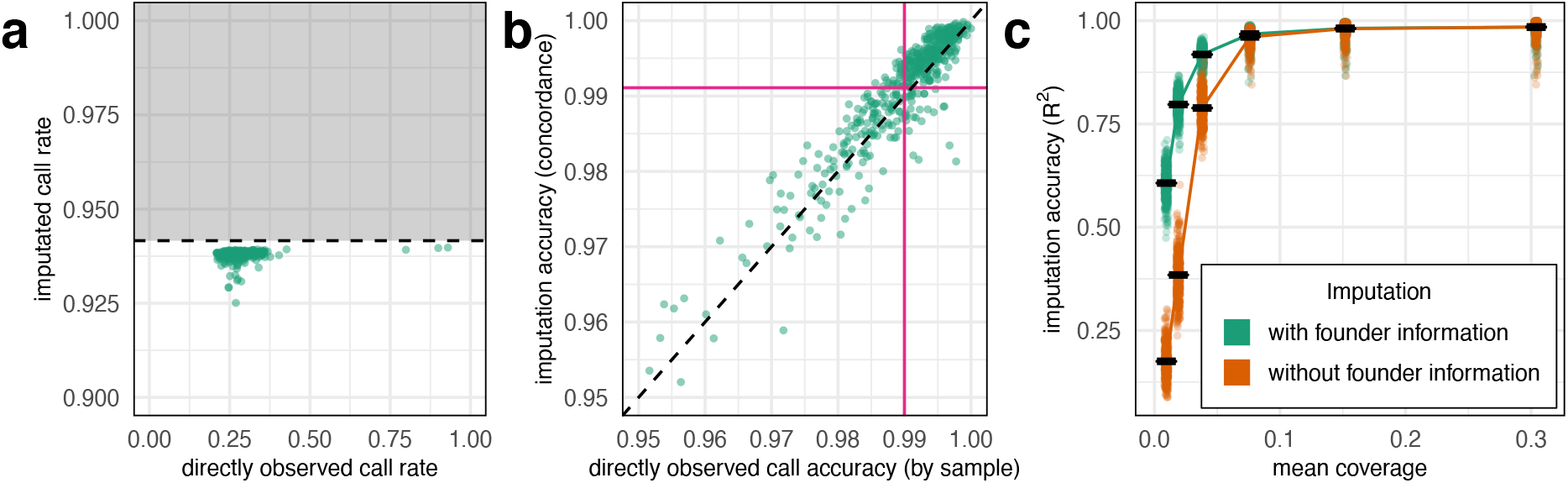
Call rate and accuracy of genotypes after imputation and after downsampling. (a) Imputed call rate (y-axis) vs direct call rate (x-axis. Only 28.1% of the 1,131,251 SNP sites can be genotyped directly from the low coverage sequence data, whereas 93.8% of sites had genotypes after imputation. 5.8% of sites (grey region and horizontal dashed line in a) were removed by quality control filters after imputation or on the unimputed ‘Un’ chromosome (0.93%). (b,c) Accuracy as evaluated at 5,747 sites that overlap with the Axiom 35k array. (c) Imputation before/after downsampling was performed with (green) and without (orange) using the genotypes of the founders as a reference panel.

### Introgressions and Segregation Bias

Several recent studies have used genomic data (e.g., SNP density[9]) to study the introgression of genetic material into hexaploid bread wheat from the secondary and tertiary gene pool[9,10,30]. We examined evidence for introgressions in previously reported locations[9,31–33] using founder coverage and non-reference allele frequency. Because we developed RILs, we were also able to examine segregation bias, which often accompanies wheat introgressions[34,35]. We found evidence for at least six introgressions covering ~1.1Gb segregating in the population, five of which showed segregation bias (Supplementary Table 3).

### Phenotypic Characterisation and QTL Mapping

We measured 47 phenotypes in replicated field trials over two years (Table 1, Supplementary Tables 4, 5, and 6), including the 10 time points at which Green Leaf Area (GLA) was measured. Of these, 25 phenotypes were collected in both years and two were also measured in smaller 1×1m nursery plots (Yellow Rust infection, YR, and Juvenile Growth Habit, JGH) to give a total of 73 phenotypic measurements. Phenotype distributions are shown in Supplementary Figure 3, showing that some RILs have more extreme phenotypes than any founder (transgressive segregation) for almost all phenotypes (RIL maximum ≥ founder maximum for 61/73 phenotypes and RIL minimum ≤ founder minimum for 68/73 phenotypes). All phenotypes have significant (p<0.05, Pearson’s correlation test) correlations with at least one other phenotype (Supplementary Figure 4).

**Table 1.** Phenotypes collected.

| ABBREVIATION | TRAIT | ABBREVIATION | TRAIT |
| --- | --- | --- | --- |
| BIS | Basal infertile spikelets | GS55 | Ear emergence date |
| EL | Ear length | GS65 | Anthesis date |
| ETA | Ear taper | GW | Grain width |
| ETS | Ear tip sterility | GY | Yield |
| EW | Ear weight | HEB | Height to ear base |
| FLA | Flag leaf angle | HET | Height to ear tip |
| FLED | Flag leaf to ear distance | HFLB | Height to flag leaf base |
| FLF | Flag leaf floppiness | JGH | Juvenile growth habit |
| FLL | Flag leaf length | LOD | Lodging |
| FLS | Flag leaf senescence | PHS | Pre-harvest sprouting |
| FLW | Flag leaf width | PIG | General pigmentation |
| GA | Grain area | SH | Spring habit |
| GL | Grain length | SPIG | Stem pigmentation |
| GLA# | Green leaf area (10 time points, Nov–Mar) | SW | Specific weight |
| GLAU | Glaucousity | TGW | Thousand grain weight |
| GPC | Grain protein content | TIS | Tip infertile spikelets |
| GR | Germination rate | TS | Total spikelets |
| GS39 | Flag leaf emergence date | YR | Yellow rust infection |

From the 1.07M imputed SNPs, we selected a subset of 55,067 pruned by linkage disequilibrium (LD). Using genome-wide association scans (GWAS) on both SNP and founder haplotype data, we mapped 136 Quantitative Trait Loci (QTLs) across the 73 phenotype/year combinations that were genome-wide significant at the 5% level. Many QTLs overlapped for different phenotypes, clustering into 42 distinct genome locations. For 25 phenotypes that were measured in both years, we found 48 QTLs in year 1 and 49 QTLs in year 2, of which 28 were mapped to the same location and were genome-wide significant in both years. For example, in replicated trials lacking fungicide treatment we mapped yellow rust (*Puccinia striiformis*) susceptibility to four QTLs in year 2 (on chromosomes 2A[31,36], 2B[37], 3B, and 6A), of which three were also mapped in year 1 (2A, 3B, and 6A); only one (6A) was also mapped in trials treated with fungicide. 126/136 QTLs at 40/42 genomic locations were mapped using SNP-based associations, whereas 87/136 QTLs at 30/42 genomic locations were mapped using haplotype-based association tests. That is, 10 QTLs and two genomic locations were only identified from haplotype-based association whereas 49 QTLs and 12 genomic locations were only identified from SNP-based association. This is consistent with the limited gene-level haplotypic diversity observed among the founders.

Figure 4b summarises the 40 loci with genome-wide significant SNP-based associations. We were able to assign 21 of these, including most of those with the strongest effects, to previously reported QTLs. In 11 high confidence cases, candidate genes have been reported and/or validated experimentally. In other cases, QTLs contained homeologs or paralogs of these high confidence candidates, or previous studies had reported associations to a genetic map using marker data, but not firmly anchored these locations on the reference genome assembly (low confidence co-localisation, n=10). We checked six high confidence candidate loci with annotated reference genome locations (*RHT-1*[38], *RHT-2*[39], *WAPO-A1*[40], *ALI-1*[41]*, TaMyb10-B1*[42], *Yr7*/*Yr5*/*YrSP*[37], *PPD-D1*[43]), all of which were within our mapping intervals. We created a genotype-phenotype map for community use by placing all QTLs on the physical map (Supplementary Table 7) to a median interval of 9.2Mb.

**Figure 4.**
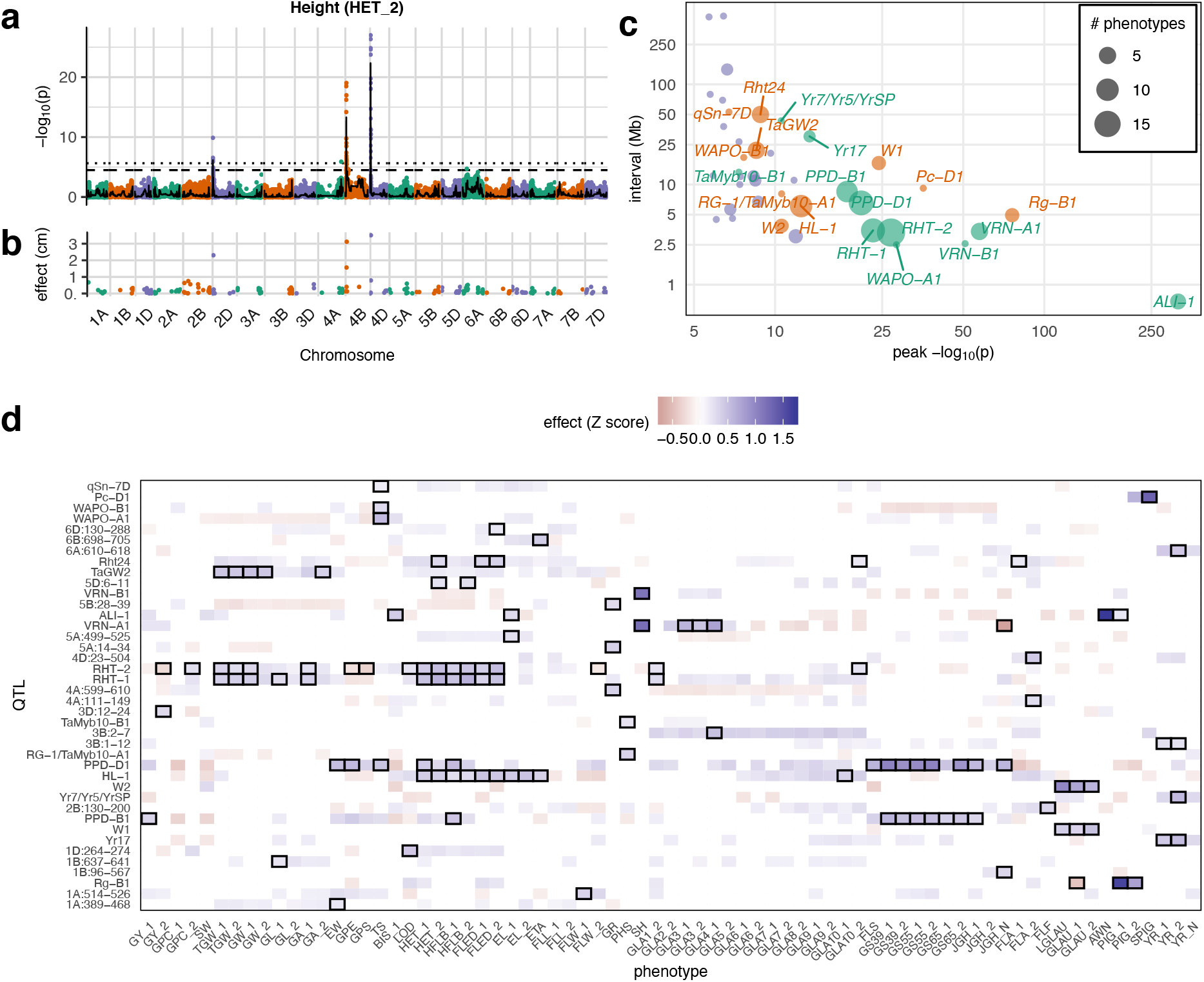
Genotype-phenotype associations. (a) Exemplar Manhattan plot of the genome-wide −log_10_ p values of association (logP) between the height to ear tip phenotype from year 2 (HET_2) and 55,067 LD-pruned SNP dosages (dots) or founder haplotype dosages (line). The horizontal lines show the 5% genome-wide significance thresholds for SNPs (dotted) and haplotypes (dashed). (b) The 193 non-zero estimated LASSO SNP effects for HET_2. (c) The 40 genomic locations where genome-wide significant SNP associations were found for at least one phenotype, classified by effect size (logP; x-axis) and genomic interval width (Mb; y-axis). Each circle represents one locus, and its size shows the number overlapping QTLs; the smallest interval width and p value is shown where there are multiple overlapping phenotype associations. Labels indicate QTLs that colocalise with previously described QTLs or candidate genes; green indicates high-confidence colocalization (n=11) and purple low-confidence colocalization (n=10). (d) Pleiotropy across 40 loci: those loci without names are labelled by chromosome and position in Mb) and 73 phenotypes. Shades indicates the significant (p<0.05) locus phenotypic effects expressed as the number of standard deviations (Z-score). Genome-wide significant QTLs are highlighted with boxes.

Most loci with strong effect co-localise with previously reported QTLs. Some large effects are commonly associated with adaptation of the founders to the geographic and temporal range they sample. For example, the early flowering allele at the photoperiod locus *PPD-D1* carried by the founder Soissons is favoured in southern Europe to avoid the summer drought[44]. The modern semi-dwarfing alleles at *RHT-B1* or *RHT-D1* that have been favoured globally since the Green Revolution[45] are absent from founders Banco, Bersee, Copain, Flamingo, Holdfast, Kloka, Spark, Steadfast and Stetson.

To examine the pleiotropic effects of the relatively few genome-wide significant QTLs, we took the most strongly associated SNP at each locus and then tested for associations with all other phenotypes, requiring a lower threshold for evidence of association (p<0.05) than was initially used to establish genome-wide significance. The results are visualised in Figure 4d, which shows that loci significant for one phenotype are also common to other phenotypes, consistent with extensive pleiotropy and the shared genetic control of correlated phenotypes.

### Gene Deletions

Our analysis of SNP variation ignored sites that could not be called reliably in all 16 founders, possibly due to whole-gene deletions relative to the reference genome. We obtained no coverage from at least one founder at 8,019 (7.2%) of genic regions and 1,095 (1.1%) of promoter-gene pairs, suggesting possible structural variations (Supplementary Figure 1). Based on the deviation in gene coverage from that expected given the mean coverage for the founder, we computed a quantitative gene deletion score (GDS) for each gene and founder and imputed the scores into the RILs using the founder ancestry mosaics. We tested the association between each GDS and each phenotype in order to identify potential causal deletions. Across 27/73 phenotypes we found 30 GDS associations with p-values <10^−6^ (Supplementary Table 8). Significant associations almost always occurred within QTLs previously mapped by SNP association, so this analysis only identified candidate genes with deletion status consistent with the pattern of action across the founders of a QTL. Of these, at 10 loci the peak GDS logP association was at least 90% of the peak SNP logP. Thus most QTLs are not likely to be caused by gene deletions. However, the GDS is based on empirical read coverage, and so is likely to be affected by stochastic experimental variations hence it is possible that the association at a true causal GDS might appear weaker than that of a tagging SNP. A further caveat is that deletions are always inferred relative to the reference genome of Chinese Spring, such that insertions or functional genes missing from the reference genome annotation will not be captured.

### Genomic Prediction

We next performed phenotypic prediction all 55,067 tagging SNPs, to predict the potential for genetic improvement within the NDM. We trained genomic prediction models using three shrinkage methods: ridge regression (RR), least absolute shrinkage and selection operator (LASSO) and Elastic Nets (EN), using 50-fold cross-validation with randomly-selected training sets comprising 90% of RILs and test sets of the remaining 10%. LASSO and EN had almost identical prediction accuracies but EN included on average 26% more SNPs than LASSO (Supplementary Figure 5). Accordingly, we only report the LASSO results. LASSO prediction accuracies for all traits are shown in Figure 5b, alongside the proportion of heritable variation explained by QTLs (Figure 5a). Across traits, LASSO had higher average prediction accuracy than RR (Figure 5c), particularly for phenotypes where a larger fraction of variation can be explained by genome-wide significant QTLs (Figure 5d), as expected for a model selection method. LASSO prediction accuracies (correlation coefficients) varied from 0.13-1 (mean 0.43) across phenotypes, using models with 1-465 SNPs (mean 155 SNPs). The number of SNPs in the LASSO model is higher for phenotypes where the overall heritability estimate greatly exceeds the fraction of variation that can be explained by genome-wide significant QTLs (Figure 5e).

**Figure 5.**
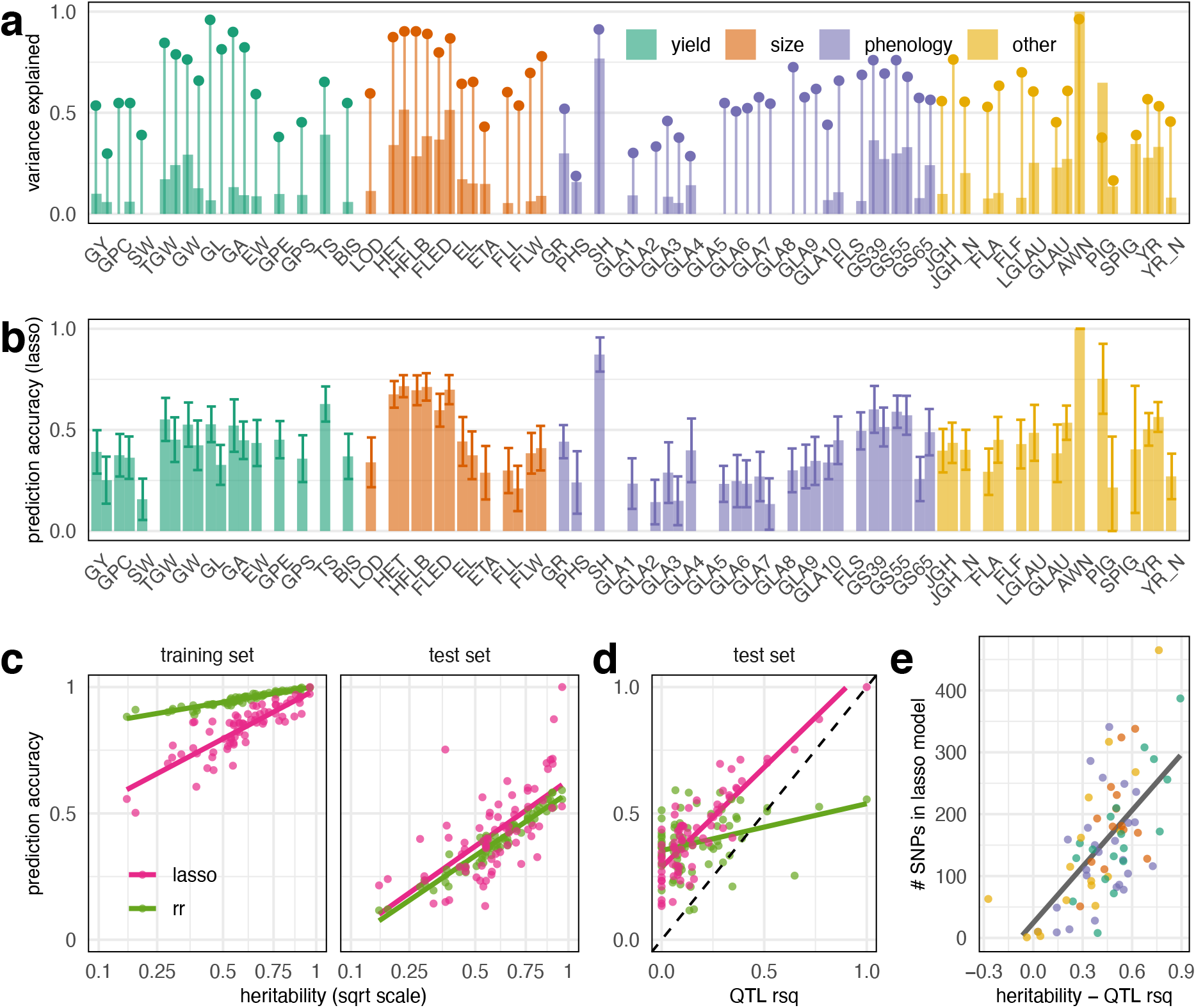
Genetic architectures of 73 trait/year combinations (47 distinct traits) as revealed by QTL mapping and genomic prediction. (a) Phenotypic variation explained by all genome-wide significant QTLs (thick bars) and by the full SNP-based genetic relationship matrix (heritability, thin bars and dots). Phenotypes measured in year 1 and year 2 are paired, shifted to the left and right, respectively. (b) LASSO prediction accuracy (correlation coefficients) across 50-fold cross validation; error bars show sds. (c) Prediction accuracy correlations (y-axis) and sqrt(heritability) (x-axis) and in the test and training sets under ridge regression (rr) and LASSO genomic prediction models. Prediction into the test set is generally higher with LASSO, especially for traits where more variation is explained by genome-wide significant QTLs (d). (e) LASSO models usually include more SNPs when more heritable variation is unaccounted by genome-wide significant QTLs (x-axis is difference between heritability and QTL R^2^).

Out-of-sample test set prediction confirms that polygenic LASSO SNPs have predictive power and are therefore likely to be tagging genetic variants affecting phenotypic variation. Most phenotypes are polygenic; their prediction models exhibited a mixture of a few large effect and many smaller effect loci. A typical example (for height) of the 193 non-zero LASSO SNP effects is shown in Figure 4c. In contrast, the Mendelian AWN phenotype is fully explained and predicted using a single genome-wide significant QTL.

The reduced accuracy of RR compared to the LASSO is expected in the absence of significant population structure. There will be reduced variation in kinship among RILs compared to the wider germplasm from which the founders are usually selected. Much of the prediction accuracy of RR results from exploiting kinship rather than from tagging causative variants[46] so there is less opportunity for high prediction accuracy in MAGIC populations. In these circumstances, a feature selection method such as the LASSO can more accurately identify and tag haplotypes contributing to trait variation and give greater prediction accuracies. The LASSO also accurately predicts traits determined predominantly by a few QTL of large effects, in which circumstances RR performs poorly (Figure 5d). The LASSO is therefore better for genomic prediction in MAGIC.

We used these genomic prediction models to explore the potential for selection in a much larger simulated population of 20,160 MAGIC RILs, 40 times larger than the real population. These were created by permuting the founder identities in the founder genome mosaics inferred in the real RILs, preserving linkage through the genetic map. Phenotypes were predicted for the test set of real RILs (10% of all lines) and in the simulated RILs for all 50 prediction models (training/test set jackknife resamples). Figure 6a shows, for two example phenotypes, that the distribution of predicted phenotypes is almost identical in the real (test set) and simulated RILs. As expected, the most extreme predicted values (maximum and minimum) in the simulated RILs exceed than those in the real dataset because novel allelic combinations are generated in the larger simulated population. However, the average improvement in extrema between the test set and simulated phenotype predictions is only −0.5 (for the minimum) and +0.68 standard deviations (for the maximum). This is in line with extreme-value distribution theory and shows that blind-breeding a very large population in the hope of generating novel combinations of beneficial alleles is inefficient.

**Figure 6.**
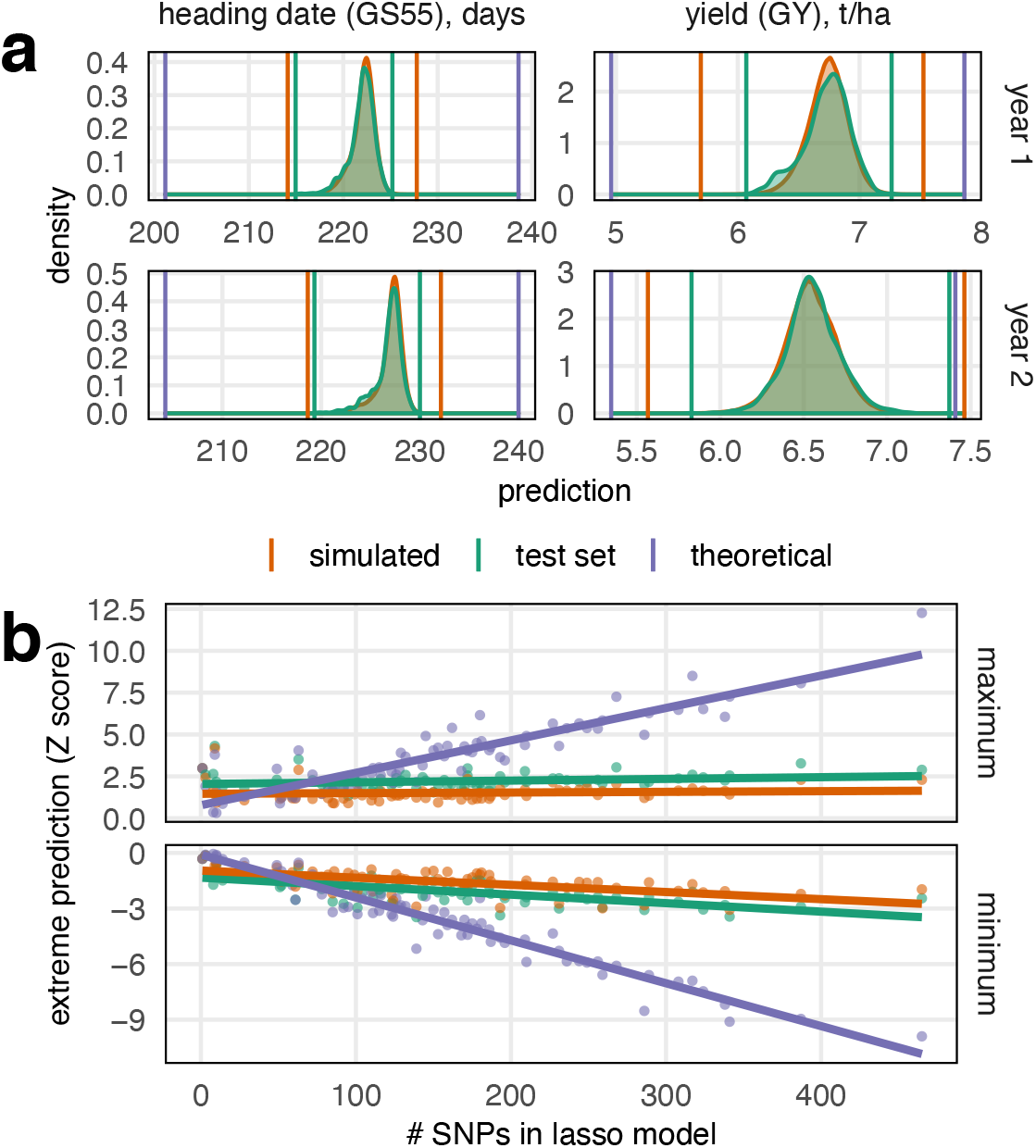
Predicted potential for phenotypic change. (a) We predicted the phenotypes of real MAGIC recombinant inbred lines, RILs (green distribution), and a large population of 20,160 simulated MAGIC RILs (orange distribution). These distributions largely overlap but more combinations are made in the simulated dataset such the the extreme values are more extreme. Nevertheless, the highest/lowest phenotypic prediction in the simulated population of 20,000 is generally only ~0.5 standard deviations higher/lower than the trait predictions in the real dataset of 504 lines. Upper graphs: predictions based on year 1 phenotype, lower graphs predictions based on year 2 phenotype (b). We also estimated the extemes of the phenotype predictions that are possible given the full lasso genomic prediction models (purple line in a). Large deviations from the current population mean are predicted to be possible but only through the fixation of a large number of loci, with less potential for change predicted at less-highly polygenic traits.

Next, we predicted the theoretical extreme phenotypic values that it is possible to create from segregating variation if unlimited recombination were possible. That is, we computed the phenotypic prediction in an imaginary line that carries all the alleles predicted to increase/decrease each phenotype. For this exercise, we trained the prediction models on the full set of 504 RILs so they differ slightly from those used to predict phenotypes in the test set. In the test set and simulated RILs, the predicted phenotypic extremes generally reflect the population size, which determines the probability that a single line happens to sample many alleles with positive/negative effects. However, Figure 6b shows that the theoretical maximum/minimum phenotypic prediction is linearly related to the complexity of the LASSO model (i.e. the number of non-zero SNP coefficients in the model). This suggests that hundreds of loci would need to be selected over multiple generations to generate any large phenotypic shifts, in line with the decades of breeding that has been required to produce genetic gain.

For essentially all crops where yield and yield quality are high priority traits, a trade-off is evident between these two phenotypes and this is recognised as a longstanding problem in wheat. Thus, identifying opportunities to break this trade-off is important[47,48]. We estimate that yield has increased by 0.021 t ha^−1^ year^−1^ based on a regression of average yield on founder release year (p=0.006, n=16, R^2^=0.43). The highest yields measured in founders and RILs exceeds the maximum predicted yield from the genomic prediction models (Figure 6) due to shrinkage in estimating SNP effects. However, high grain yield (GY) is correlated with low grain protein content (GPC) among the founders (Pearson’s correlation coefficient −0.94, p<0.001, n=16), Figure 7. Founder genetic material is reshuffled without selection in the RILs, but the GY-GPC relationship continues (correlation −0.77, p<0.001, n=504), suggesting some pleiotropy in the underlying genetic effects. To investigate the segregating genetic variation that may be available to break this trade-off, we analysed the deviation from the trend (PYD: distance from symmetrical Thiel-Sen regression between GPC and GY, after Z-score normalisation). The heritability for PYD was 0.41 in year one and 0.25 in year two and could be predicted with accuracy 0.26 (sd 0.11) in year one and 0.13 (sd 0.11) in year two. These estimates are lower than those for GY and GPC analysed separately (GY heritability 0.54 and 0.30, prediction accuracy 0.39 and 0.25; GPC heritability 0.55 and 0.55, prediction accuracy 0.375 and 0.36, Figure 5. PYD of the founders did not correlate with release date, but these results suggest modest potential to break the yield-protein trade off, requiring strong and targeted breeding effort[47,48].

**Figure 7.**
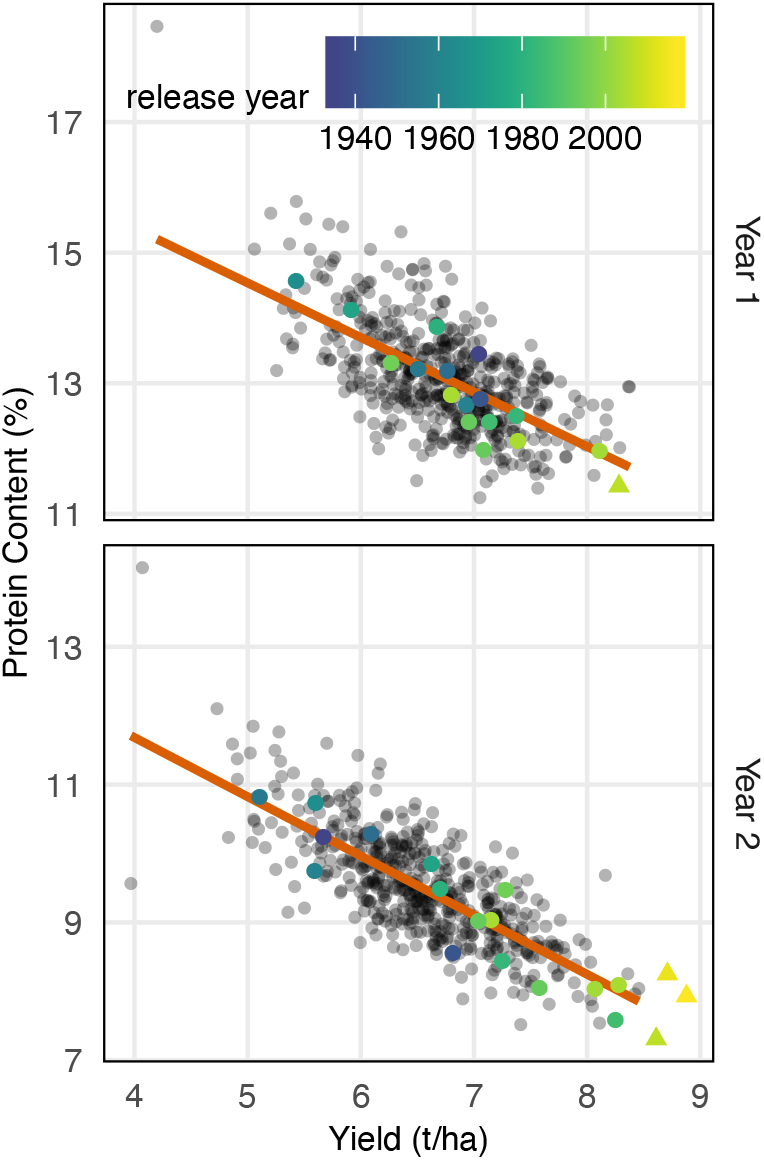
Negative trade-off across two years between grain yield (GY; x-axis) and grain protein content (GPC; y-axis) in 504 NIAB DIVERSE MAGIC RILs, 16 founders, and 3 more recently developed varieties (triangles, only one measured in year 1).

## Discussion

We report five main findings. First, imputation from low coverage WGS is a cost-effective and straightforward genotyping strategy for crops, at least in multiparental populations. Despite its large, repetitive and hexaploid genome, wheat genotypes can be reliably imputed from WGS with average per-sample coverage in the range of 0.075x-0.3x and without the use of reference panels[26]. Thus there is no absolute requirement to even know the identities of, let alone sequence, the population founders, although this may be desirable for other purposes such as pan-genome assembly and re-annotation[18,49]. In this study, we were able to impute genotypes and founder haplotypes at >1M SNP sites in >500 NDM RILs, which proved ample for genetic mapping and genomic prediction.

Second, based on SNPs called from exome capture, no more than three haplotypes segregate at most genes in commercial NW European bread wheats released since 1935. There appears to be little or no variation at about a quarter of genes on the A and B subgenomes, and at about half on the D subgenome. Complete re-assembly and re-annotation of the 16 founders of the NDM would yield more complete insights into the extent and impact of coding variation. Limits on haplotypic variation are probably the result of historical selection and population bottlenecks that reduced the effective population size before the onset of intensive breeding programmes[12,50], as well as the close relatedness among breeding materials in more recent wheat pedigrees[51]. However, it appears that the low overall level of genetic diversity has not been further reduced during the 20^th^ Century[52,53].

Third, as a consequence, most QTLs are accounted for by bi-allelic SNPs rather than haplotype differences. For comparison, about 40% of QTLs identified in a multi-founder population of rats were attributed to multi-allelic/haplotypic effects[54]. Furthermore, most genome-wide significant QTLs had pleiotropic effects. Extensive pleiotropy suggests that naïve selection on one phenotype is likely to induce correlated responses in other phenotypes. In particular, we found improved yields in recent varieties has come at the cost of a decline in protein content (Figure 7; increasing yield by one t/ha reduces protein content by about 1%). Despite reshuffling haplotypes without selection, this trade-off continues in the NDM, which indicates directed selection would be required to break the yield-quality trade-off, potentially creating varieties with improved nitrogen use efficiency[55,56].

Fourth, across 47 phenotypes, we found a wide range of underlying genetic architectures. For traits such as awns, pigmentation, spring habit and yellow rust resistance, almost all of the heritable phenotypic variance could be explained by one to four genome-wide significant QTLs (Figure 5). In other cases, a few loci with large phenotypic effects were accompanied by dozens of loci with smaller effects on traits such as flowering time and height (Figure 4a). The loci with very large effects have mostly been reported before (Figure 4b) because we recapitulate key historical steps such as the introduction of photoperiod sensitive and semi-dwarfing alleles from Japan[44,45]. Traits such as yield were polygenic with the majority of heritable variation coming from many loci of smaller effect (Figure 5).

Fifth, our genomic prediction models suggest that hundreds of loci will need to be selected and fixed to achieve large phenotypic changes in polygenic traits in the future (Figure 6B). We achieved reasonable prediction accuracy with modest numbers of SNPs; the mean out-of-sample prediction accuracy was 0.43, using on average only 155 SNPs per phenotype, out-performing ridge-regression which considers all markers simultaneously. Other crop and livestock studies have also found very sparse markers can be sufficient for useful genomic prediction[11,57,58]. Here, rather than using low marker densities, we trained models that select a few hundreds of markers from ~55k tagging SNPs. In part, this sparsity is a consequence of the design and construction of MAGIC populations, eliminating rare alleles and creating blocks of markers that can be easily tagged in prediction models[15]. These factors may be responsible for the use of far fewer markers than used to generate polygenic prediction scores in humans[59], where there is a long tail of rare variation and less linkage disequilibrium.

Our results suggest that dramatic genetic improvement over 70 years of breeding has largely been achieved through the fine shuffling of a low number of haplotypes to recombine polygenic alleles of small effect, combined with the introduction of alien introgressions from wide crosses. The introgression of large genomic segments from related species has most commonly been for sources of resistance to specific diseases[9,33,34]. Breeders now have a choice whether to continue with the same strategy, i.e. selecting from within existing variation and introgressing selected exotic alleles, or to ambitiously expand the pool of available haplotype diversity genomewide.

## Methods

### NDM Population Creation

The 16 NDM founders were chosen to capture the greatest genetic diversity using PowerMarker genetic analysis software[60]. They were chosen from 94 NW European wheats released in the UK that were genotyped with 546 DArT and 61 SSR markers; the full panel also included 96 US and 50 Australian varieties, which were excluded based on STRUCTURE analysis[61]. The founder selection process was run iteratively with the varieties ‘Robigus’ and ‘Soissons’ first fixed to be included to coincide with the founders of the 8-founder NIAB Elite MAGIC population[62]. Then the most frequently selected additional 4, then 9, and 12 varieties were fixed in multiple iterative selection runs and finally the most frequently selected 16 were chosen. Seed for the founding varieties was sourced from the John Innes Centre Germplasm Resource Unit (GRU http://www.jic.ac.uk/germplasm/).

These founders were inter-crossed in a balanced funnel crossing scheme, based on a Latin square field trial design, over four generations to create 16-way crosses with all the founders equally represented in their pedigree. First, all 120 possible 2-way crosses between founders were made in a half diallel scheme. Two-way plants were then crossed in 60 4-way combinations. Multiple plants from each family were used in crossing from 2-way onwards, in order to maintain maximum founder allelic diversity within the population. 30 crossing combinations were made between 4-way plants to create 8-way crosses, making between five and eight replicate crosses per combination using different plants. These were intercrossed in 15 combinations to create balanced 16-way crosses, with each combination replicated between six and fifteen times using different 8-way plants. This resulted in 174 16-way plants from which one to sixteen inbred lines per 16-way family were made through single seed descent (SSD). 596 RILs were advanced to the F_7_ stage when seed for phenotyping was multiplied in 1×1m nursery plots. Supplementary Table 9 gives details the number of plants involved in each cross and Figure 2a shows the pedigree for the 504 RILs used in our main analysis only.

### Phenotyping

RILs from the population were phenotyped in field trials over multiple environments near Cambridge, UK. Yield trials were conducted in the growing seasons 2016-2017 and 2017-2018, hereafter year 1 and year 2 (phenotype suffix codes _1 and _2). Information on location, soil type, key dates and inputs for both years are given in Supplementary Table 4. Yield plot dimensions were 2m wide and 4m long and plots were sown at a density aiming to achieve 300 plants m^−2^. In year 1, 596 lines were included in two replicates, the sixteen founders in four replicates and the commercial control variety ‘KWS Santiago’ in 24 replicates in a randomised nested block design with 16 main blocks of 80 adjacent plots which comprised each row in the trial and eight sub-blocks of ten plots nested within each main block. In year 2 trials, 596 lines and the 16 founders were included in two and four replicates respectively but three control varieties (‘KWS Santiago’, ‘Skyfall’ and ‘Shabras’) were all included in four replicates. Plots were again randomised in a nested block design but including additional plots making a larger trial, consisting of 20 main blocks of 115 adjacent plots, which comprised each row, and 23 sub-blocks of five plots nested within each main block.

Disease observation trials (DOTs) were conducted near Cambridge, UK in the same years as the yield trials to assess resistance to crop diseases. These plots consisted of two 1.2m length rows, treated with no fungicide but otherwise standard inputs. Due to local conditions, DOTs were considered to have natural high pressure of yellow rust (*Puccinia striiformis* f.sp. *tritici*). In both years, DOTs included two replicates of 596 RILs, four replicates of the 16 founders and 68 additional replicates of the susceptible founder ‘Robigus’. Trial designs included two main blocks of 660 plots, with 11 sub-blocks of 60 plots nested within main blocks. All trial designs for both yield and disease observation trials were made using the package ‘blocksdesign’ in R. Phenotyping of some traits was also carried out in 1×1m seed nursery plots where lines were not replicated but the founders were in three replicates and randomised across the nurseries (phenotype code _N).

A wide range of traits were phenotyped across the field trials, including traits for crop developmental morphology, phenology, plant stature and canopy architecture, yield and yield components such as spike and grain morphology, disease resistance, pigmentation, plant glaucosity, indications of stress response, lodging, grain protein content and vernalisation requirement. A summary of these traits and abbreviations are presented in Table 1 and details of phenotyping methods are listed in Supplementary Table 5.

### Trials Analysis

Adjusted phenotype values were calculated as Best Linear Unbiased Estimates (BLUEs) for each trait separately for each trial year using mixed effects models with ASRemL[63]. Genotype was considered a fixed effect whilst experimental blocking structure as well as other covariates such as harvesting day, where relevant, were included as random effects. Spatial models including first- and second-order auto-regressive spatial models were also used. Model simplification was carried out where models with all possible combinations of random effect terms and spatial terms for row and column were run and the best fitting model was chosen based on Akiake Index Criteria (AIC). Model residuals were visually checked for normality and equal variance to fitted values distribution. Best Linear Unbiased Estimates (BLUEs) for all phenotypes for the 16 founders and for the 504 RILs used in our main analysis (see below) are provided in Supplementary Table 6. We used symmetrical Thiel-Sen regression (implemented in the ‘deming’ R package) after phenotype normalisation to characterise the relationship between protein content (GPC) and yield (GY). The Protein-Yield Deviation (PYD) phenotype is calculated as the Euclidian distance from this regression line.

### Genotyping Array Data

All DNA extraction was performed using the Qiagen DNeasy Plant Kit on leaf tissue samples taken from emerging leaves of seedlings. First, genotyping was performed at the Bristol Genomics Facility using the Axiom 35k wheat breeders’ array[12]. Initially, two 384-sample plates were genotyped. Seed from the plants used as founders were genotyped on each plate (32 samples) along with extra seed from the original varietal seed stock used (28 samples) and seed from founders propagated to 2017 (16 samples). In addition, 596 RILs were genotyped after 5 generations of selfing (F_6_). To account for genotyping failures and to ensure the accuracy of sample labels, 150 RILs were re-genotyped in the F_7_ generation along with a further replicate of each founder.

Genotype calling was performed using the Affymetrix Power Tools (v1.19) and SNPolisher R packages, following the recommended Axiom analysis pipeline. All samples except two-way crosses were given the standard inbreeding penalty, 4, which penalises calling heterozygous genotypes. Four samples failed the ‘dish quality control’ threshold (0.82) and a further 28 samples with call rates were below 97% were excluded. Marker classifications were performed using “ps-classification”, and ps-classification-Supplementary” functions with options --species-type polyploid --hom-ro false. All calls were adjusted using the standard 0.025 confidence threshold using the Ps_CallAdjust function.

Samples were compared to one another using the 14,935 markers classified as ‘PolyHighResolution’ only. Overall, 46 RIL pairs were found to be >92% similar (mean 98.5% genotype similarity), where all other comparisons between MAGIC lines were, at most, 84% similar (mean 67.8%). These apparently duplicated genotypes could indicate genotyping, labelling, or propagation errors so only one RIL from each pair was used for sequencing (550 RILs). To ensure pedigree accuracy, we chose the RIL in each pair that was genotypically most similar to other RILs derived from the same 16-way cross (i.e. in the same family).

### Sequencing Data

For whole genome sequencing, DNA was extracted from 550 RILs at the F_7_ generation. DNA for RILs that failed quality control were extracted again at the F_8_ generation (n=50). Sequencing and library preparation was performed at Novogene, where libraries were generated from 1.0μg DNA per sample using the NEBNext DNA Library Prep Kit. Sequencing was performed on a NovaSeq 6000 instrument (Illumina) to get at least 6Gb of raw sequence data (2×150bp paired end reads) per sample. One founder (Holdfast) was sequenced to 15.8x coverage using the same method.

The other founders were sequenced after capture using two recently designed probe sets targeting promoter and genic regions, respectively[21]. Capture was performed at the Earlham Institute following the SeqCap EZ Library SR v5.1 protocol (Roche NimbleGen Inc., Madison, WI, USA) with 1μg of genomic DNA sheared to 300bp[21]. Four captures were performed using 8 samples per set (2x promoter captures and 2x genic captures). Samples for the founder Stetson were included on all four capture experiments so roughly double the sequence data was obtained for this variety (Supplementary Table 1). Sequencing with 2×150bp reads was performed at the Earlham Institute on a NovaSeq 6000 instrument (Illumina) with 16 promoter capture libraries on one lane and 16 genic capture libraries on another lane.

### Variant Calls and Imputation

All reads were aligned to the bread wheat reference genome from cv. Chinese Spring (RefSeq v1.0)[27] using bwa-mem (version 0.7.12)[64] and sorted using samtools (version 1.3.1)[65], which was also used to calculate coverage. For compatibility with the bam file format, we split each chromosome in the reference genome at the halfway point before alignment. We called variants from the founder sequences within the high confidence gene, promoter and 5’ UTR regions targeted by the capture probes[21] using GATK (version 4.0.8.0)[66] HaplotypeCaller and GenotypeGVCFs (options --interval-padding 100 --minimum-mapping-quality 30). We used vcftools (version 0.1.15) to include only bi-allelic single nucleotide polymorphisms (SNPs) with average coverage depth between 5 and 60 (all per sample coverages between 2 and 120) and no missing calls. We also filtered with bcftools (version 1.2)[67] using standard quality control options --exclude’QD<2 || FS>60.0 || MQRankSum<−12.5 || ReadPosRankSum<−8.0 || SOR>3.0 || MQ<40’. This left 1.78M SNPs, of which we only use the 1.13M sites with no heterozygous calls (--genotype ^het option) for our main analyses.

We first called genotypes in the RILs at these 1.13M SNP sites directly using GATK HaplotypeCaller in GENOTYPE-GIVEN-ALLELES mode, using the same options as above. We assessed the concordance between array genotypes and these direct calls (AD) at overlapping sites (see below). For 10 RILs, the directly called sequencing variants most closely matched genotyping array data for a different line than expected. These were excluded because the source of the discrepancy (sequence data or array data) cannot be established. The concordance between our genotyping array data and direct calls (AD) was below 95% for a further 36 RILs, which were excluded (mean AD 84.7% for removed lines), leaving 504 RILs. We estimated heterozygosity in these 504 RILs using only genotypes called from at least four reads. Of 2.6M such genotype calls, only 0.67% were called as heterozygotes.

We imputed genotypes at the 1.13M SNP sites using the alignments and STITCH software (version 1.5.7)[26]. Because alignments were to a reference genome with chromosomes split in half, we first ran STITCH with the generateInputOnly option, and then joined the input files for each chromosome half before imputation. For all runs, we used the parameters nGen=3, minRate=0.001, bqFilter=30, method=‘diploid-inbred’ and then filtered all sites with an info score below 0.4, minor allele frequency below 2.5%, or missingness above 10%. For our main analysis, we used the genotype calls in the founders as a reference panel and outputted the estimated ancestry dosages of each founder at each position in each RIL using the outputHaplotypeProbabilities and output_haplotype_dosages options. When using the founders as a reference panel, we removed options that estimate and update the haplotypes in the population (shuffleHaplotypeIterations, reference_shuffleHaplotypeIterations, refillIterations). To test accuracy when reference panels aren’t available, we re-ran imputation without the founder haplotypes, using 40 iterations to estimate the haplotype space and recombination mosaics. We also used the downsampleFraction option to randomly sample a fraction of alignments with/without using the founder reference panel. Finally, we tested imputation accuracy (without a reference panel), when fewer than sixteen haplotypes were assumed to segregate in the population by varying the K parameter (Supplementary Figure 2).

### Genotype Comparisons

For comparison against the sequencing dataset, we used all genotyping array markers. Replicates of founders and MAGIC RILs (where available) were used to make a consensus call where the most common genotype across replicates was taken as the consensus and only retained when more than 50% of the non-missing calls were in agreement. In addition, markers where one homozygous genotype was missing from all RILs were converted such that all heterozygous calls were assumed to be in the missing homozygous class. The failure to detect a homozygous class is likely to be a result of polyploidy, which can reduce differentiation between the three genotype classes and make them hard to distinguish. Finally, to get plausible physical positions for the genotyped markers, BLASTn v2.2.30[25] was used to compare the 75bp probe sequences (cerealsdb.uk.net)[12] against the reference genome[27]. When matching the SNP array data with the sequenced SNPs, array sites were excluded if there had missing or heterozygous founder calls or if the genotypes and targeted SNP alleles did not match the founder sequence data. We found 5,877 sites that overlapped between the genotyping array data and the sequencing data (Supplementary Table 2).

To compare against global wheat diversity, we called founder genotypes at 113,457 genotyping array sites that were polymorphic among 4,506 diverse global wheat accessions[13]. We called genotypes from alignments with mapping quality scores of at least 30 using GATK HaplotypeCaller in EMIT_ALL_SITES mode with the –emit-ref-confidence BP_RESOLUTION option, providing a bed file of the 113,139 genotyping array sites[13]. We only considered sites where genotypes could be called in all 16 founders (n=56,063). We used genotyping array calls for cv. Chinese Spring to determine reference/non-reference alleles on the genotyping array, ignoring sites called as heterozygous (n=109) or missing (n=306) in Chinese Spring. Seven of the MAGIC founders were also present in the global genotype set (Brigadier, Copain, Maris Fundin, Soissons, Spark, Steadfast, Stetson)^7^. The average concordance of the global genotype calls and our sequencing calls for these founders was 94.3% (sd 0.63%). We excluded 5,491 (9.8%) sites that had mismatches across these founders, many of which are likely to reflect differences in the underlying genetic variation picked up by the different genotyping technologies. Two other founder variety names were in the genotyping array dataset^7^ (Banco and Holdfast) but the genotyping calls did not match (concordances 74.2% and 71.4%, respectively), which may reflect differences in the seed stock used.

### Founder Haplotype Diversity

First, we used the SNPs called within each promoter-gene pair to estimate haplotypic diversity of the founders. We calculated absolute (Manhattan) pairwise genetic distances between founders at each site and then used complete linkage clustering to define haplotypic groups using dist and hclust functions implemented in R statistical software (version 3.6.0)[68]. This was repeated using different similarity thresholds to define haplotypes. Second, we determined haplotype breakpoints using a dynamic programming algorithm. For each pairwise founder combination, our algorithm calculates a mosaic of genotypic similarity/dissimilarity akin to the Viterbi path from a hidden Markov model. Genotype matches and mismatches are allocated a score (1 by default). To prevent excessive switching between states, there is also a ‘transition penalty’ for inferring a change between matching and mismatching states. Based on their pairwise matching/mismatching states, we then infer the total number of haplotypes inferred at each site. We repeat this procedure with different parameter choices (Supplementary Figure 2).

### Genetic Mapping and Heritability

For mapping, we used the full set of 1,065,185 high-quality SNP sites called in 504 RILs after imputation and quality control filters. We also selected a subset of 55,067 SNPs such that every other SNP was tagged at R^2^>0.99 by a member of the subset using PLINK (version 1.90) with option --indep-pairwise 500 10 0.99. These tagging SNPs were used to calculate the genetic relationship matrix ***K*** = ***GG’***/*p*. The phenotypic variance-covariance matrix for a given vector y of standardised phenotype values was modelled as 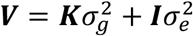. where 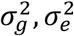. are the additive genetic and environmental variance components, estimated by maximum-likelihood[69]. The heritability of a trait was defined as 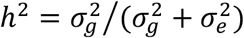. The matrix square root of the variance matrix was calculated by eigendecomposition of ***V*** as ***A***^2^ = ***V***, and the mixed model transformation of the data performed i.e. ***y*** → ***A***^−1^***y***, ***G*** → ***A***^−1^***G***, ***V*** → ***I*** to remove the inflationary effects of unequal relatedness on genetic associations before association mapping.

We performed association tests at the level of both SNPs and founder haplotypes using R statistical software (version 3.6.0)[68]. Initially, we tested the null hypothesis of no association at each SNP site in the tagging set (~55k sites). We then determined genome-wide thresholds for statistical significance using 1,000 permutations on the transformed phenotypes. If any association exceeded the 0.05 threshold (smaller p value than found across at least 950 phenotypic permutations), then we repeated the association test at all of the ~1.1M SNPs on the chromosome with the strongest association signal (lowest p value). Mapping intervals were defined to include SNPs surrounding the peak SNP, with log_10_(p) values within *d* units of *x* using *d* = max {2, 0.1*x*} where *x* is the peak log_10_(p) value. The interval for haplotype-based tests includes the range of sites that have log_10_(p) values within *d* units of *x*. SNP-based intervals were calculated using the same measure but then extended by the minimum of 5Mb or the distance to the next SNP in either direction that the same ‘strain distribution pattern’[54] as any highly-associated SNPs (SNPs with log_10_(p) values within *d* units of *x*). The ‘strain distribution pattern’ is the pattern of major/minor alleles across founders. This procedure is designed to capture the uncertainty in the positioning of relevant recombination events either side of the QTL peak. We fitted QTLs in a stepwise manor by fitting the phenotype against the most strongly associated SNP (or haplotype dosage) whenever genomewide significant QTLs were detected. The above association test procedure was then repeated using the phenotype residuals after fitting all previously identified QTLs. This allows closely-linked QTLs to be detected when they have different patterns of causal variants among RILs. Where QTL associations were found for different genotypes, they were judged to be at the same locus if they had overlapping mapping intervals and at least one matching strain distribution pattern at highly-associated SNP sites.

### Genomic Prediction

To evaluate the accuracy of trait prediction within our magic population and estimate the extent of polygenic variation beyond genomewide significant QTLs, we conducted genomic prediction across all phenotypes using three shrinkage-based methods: ridge regression (RR), Elastic Nets (EN) and least absolute shrinkage and selection operator (LASSO). We note that with appropriate choice of ridge parameter 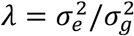, RR is equivalent to a mixed model in the sense that the RR estimated SNP effects are identical to the mixed-model Best Linear Unbiased Predictors (BLUPs)[70,71]. This explains the linear correlation between estimates of heritability and RR prediction accuracy (Figure 5c). For each method, we conducted 50 rounds of cross validation by randomly sampling 90% of the RILs (n=454) as a training set in each round to train the model, which was then used to predict the remaining 10% of RILs (n=50) - the test set. For the three methods, the model equation can be written generally as *y* = *μ* + *βG* + *ε*, where *y* is the estimated trait value, *μ* is the model intercept, *β* is the vector of SNP effects, *G* is the genotype dosage matrix, and *ε* is the residual error.

The genomic prediction models were trained using the R package glmnet[72], which estimates an optimal lambda shrinkage value for all three genomic prediction methods based on the training set. We then predicted phenotypes in the test set by multiplying all SNP coefficients estimates by their corresponding genotypes in the test set (and adding the intercept term). We report the training and test set prediction accuracy as the mean Pearson correlation coefficient of the predicted trait values and the actual phenotype values over 50 rounds of cross validation.

We used these genomic prediction models to simulate the potential for phenotypic change. First, we permuted the population founder haplotypes identities 40 times across 504 RILs and then projected the permuted founder genotypes onto the new lines. This creates new genetic combinations while retaining the genetic map and linkage found in the real population. We then used the three models trained as described above to predict phenotypes for the simulated MAGIC RILs. We further calculated the theoretical maximum and minimum phenotype values that are possible given the genomic prediction models and the variants segregating in the population. To estimate the maximum and minimum potentially achievable phenotype values, we trained new genomic prediction models using the full data set of 504 RILs for all phenotypes. We then calculated the maximum/minimum predicted phenotypes by summing the estimated effects for all positive/negative SNP coefficients.

### Gene Deletion Analysis

We examined the power of gene-level coverage variation among founders to explain phenotypic variation. In each founder *f* and at each gene feature *g*, we computed a deletion index *D*_*gf*_based on the number of reads aligning to the associated capture sequences, normalised by the overall coverage for that founder. The gene deletion score (GDS) for each MAGIC RIL *i* and feature *j* was computed as *S*_*ij*_ = ∑_*f*_ *H*_*ijf*_*D*_*jf*_, where *H*_*ijf*_ is the haplotype dosage for founder *f* in RIL *i* at gene *j*, as computed by STITCH. For each phenotype a mixed-model GWAS was performed, using the GDS in place of SNP dosages and with a genetic relationship matrix computed from the GDS (Supplementary Table 8). We also repeated the genomic prediction analysis described above by replacing the SNP genotype dosage matrix with the GDS matrix (Supplementary Figure 5).

### Introgressions

The presence of introgressions were determined using summary statistics (coverage, non-reference allele frequency in founders and RILs) calculated in 10Mb windows moved in 5Mb steps. Within introgressions, carriers have a high proportion of non-reference alleles due to the alignment of inter-specific genetic material to the bread wheat reference genome. The introgression extent was determined as the extent of 10Mb windows where all introgression carriers had a higher proportion of non-reference alleles than all non-carriers. Within these regions, we then checked the relative coverage of carriers and the extent to which the alleles of carriers are over- or under-represented among the RILs. This evidence is summarised in Supplementary Table 3. For example, the founder Maris Fundin carries a large introgression (640Mb) from *Triticum timopheevi* on chromosome 2B that inflates the total number of SNPs called on chromosome 2B, relative to the other chromosomes (Supplementary Figure 1), this introgression is substantially over-represented among RILs, as expected[34].

## Acknowledgements

The initiation of the NIAB DIVERSE MAGIC (NDV) population was funded by Biotechnology and Biological Sciences Research Council (BBSRC) grant BB/E007201/1, awarded to IM. Completion of the population, and its phenotypic and genetic analysis was funded by grants BB/M011666/1 to JC and BB/M011585/1 to RM, with some interim NDV germplasm development advancement supported by grant BB/I002561/1. Genomic prediction analyses were supported by grant BB/P024726/1 to RM. The small-plant phenotyping at IBERS was funded by BB/M011666/1 with additional support from BBSRC National Capability Grant BB/CAP1730/1. We thank Ana Sanchez and the temporary staff at NIAB who contributed to phenotypic data collection, and the NIAB Trials Team for management of field trial sites.

## Author Contributions

Project conception and funding: IM, PH (BB/E007201/1) and RM, JC (BB/M011666/1, BB/M011585/1). Project management: JC, RM, with input from MS, NF, PH, KG and AB. Population design IM, PH. Population creation: RH, NF. Seed preparation for trials, phenotyping, trials analysis and DNA extraction: NF. Small plant phenotyping: FC. Promoter-gene capture pull downs and library preparation: TB. Genetic analyses: MS with support from RM. Genomic prediction: OL. Manuscript writing: MS, OL, NF, RM, with inputs from KG, IM and JC. All authors edited and approved the manuscript.

## Availability of data and materials

The sequence datasets supporting the conclusions of this article are available in the ENA repository, under project number PRJEB39021 (temporarily embargoed), https://ebi.ac.uk/ena.

The Genotyping array genotypes for founders and MAGIC RILs are available from http://mtweb.cs.ucl.ac.uk/mus/www/MAGICdiverse/index.html in text tabular format.

The Imputed SNP genotypes and founder haplotype dosages are available from mtweb.cs.ucl.ac.uk/mus/www/MAGICdiverse/MAGIC_diverse_FILES/MAGIC_PLINK.tar.gz, and mtweb.cs.ucl.ac.uk/mus/www/MAGICdiverse/MAGIC_diverse_FILES/MAGIC_HAPLOHAPLO.tar.gz (temporary links).

The remaining datasets supporting the conclusions of this article are included within the article and its additional files.

Custom analysis scripts (mixed model and haplotype dynamic programming algorithm) are available from github.com/michaelfscott/DIVERSE_MAGIC_WHEAT.

## References

1. FAO. Crop harvested area (yield and production). Crop Harvest. area (yield Prod. GAEZ. 2014.

2. Tester M, Langridge P. Breeding technologies to increase crop production in a changing world. Science (80-). 2010;327:818–22.

3. Brancourt-Hulmel M, Doussinault G, Lecomte C, Bérard P, Le Buenec B, Trottet M. Genetic Improvement of Agronomic Traits of Winter Wheat Cultivars Released in France from 1946 to 1992. Crop Sci. 2003;43:37–45.

4. Shearman VJ, Sylvester-Bradley R, Scott RK, Foulkes MJ. Physiological processes associated with wheat yield progress in the UK. Crop Sci. 2005;45:175–85.

5. Sanchez-Garcia M, Royo C, Aparicio N, Martín-Sánchez JA, Álvaro F. Genetic improvement of bread wheat yield and associated traits in Spain during the 20th century. J Agric Sci. 2013;151:105–18.

6. Mackay I, Horwell A, Garner J, White J, McKee J, Philpott H. Reanalyses of the historical series of UK variety trials to quantify the contributions of genetic and environmental factors to trends and variability in yield over time. Theor Appl Genet. 2011;122:225–38.

7. IWGSC. Shifting the limits in wheat research and breeding using a fully annotated reference genome. Science [Internet]. 2018;361:eaar7191. Available from: http://science.sciencemag.org/content/361/6403/eaar7191

8. Adamski NM, Borrill P, Brinton J, Harrington SA, Marchal C, Bentley AR, et al. A roadmap for gene functional characterisation in crops with large genomes: Lessons from polyploid wheat. Elife. 2020;9:1–30.

9. Cheng H, Liu J, Wen J, Nie X, Xu L, Chen N, et al. Frequent intra- and inter-species introgression shapes the landscape of genetic variation in bread wheat. Genome Biol. Genome Biology; 2019;20:1–16.

10. He F, Pasam R, Shi F, Kant S, Keeble-Gagnere G, Kay P, et al. Exome sequencing highlights the role of wild-relative introgression in shaping the adaptive landscape of the wheat genome. Nat Genet [Internet]. Springer US; 2019;51:896–904. Available from: http://dx.doi.org/10.1038/s41588-019-0382-2

11. Juliana P, Poland J, Huerta-Espino J, Shrestha S, Crossa J, Crespo-Herrera L, et al. Improving grain yield, stress resilience and quality of bread wheat using large-scale genomics. Nat Genet [Internet]. Springer US; 2019;51:1530–9. Available from: http://dx.doi.org/10.1038/s41588-019-0496-6

12. Winfield MO, Allen AM, Burridge AJ, Barker GLA, Benbow HR, Wilkinson PA, et al. High-density SNP genotyping array for hexaploid wheat and its secondary and tertiary gene pool. Plant Biotechnol J. 2016;14:1195–206.

13. Balfourier F, Bouchet S, Robert S, DeOliveira R, Rimbert H, Kitt J, et al. Worldwide phylogeography and history of wheat genetic diversity. Sci Adv. 2019;5.

14. Hufford MB, Berny Mier Y Teran JC, Gepts P. Crop Biodiversity: An Unfinished Magnum Opus of Nature. Annu Rev Plant Biol. 2019;70:727–51.

15. Voss-Fels KP, Stahl A, Wittkop B, Lichthardt C, Nagler S, Rose T, et al. Breeding improves wheat productivity under contrasting agrochemical input levels. Nat Plants [Internet]. Springer US; 2019;5:706–14. Available from: http://dx.doi.org/10.1038/s41477-019-0445-5

16. Huang BE, Verbyla KL, Verbyla AP, Raghavan C, Singh VK, Gaur P, et al. MAGIC populations in crops: current status and future prospects. Theor Appl Genet. 2015;128:999–1017.

17. Cavanagh C, Morell M, Mackay I, Powell W. From mutations to MAGIC: resources for gene discovery, validation and delivery in crop plants. Curr Opin Plant Biol. 2008/02/26. 2008;11:215–21.

18. Scott MF, Ladejobi O, Amer S, Bentley AR, Biernaskie J, Boden SA, et al. Multi-parent populations in crops: a toolbox integrating genomics and genetic mapping with breeding. Heredity (Edinb). Heredity (Edinb); 2020;

19. Ladejobi O, Elderfield J, Gardner KA, Gaynor RC, Hickey J, Hibberd JM, et al. Maximizing the potential of multi-parental crop populations. Appl Transl genomics. Appl Transl Genom; 2016;11:9–17.

20. White J, Law JR, MacKay I, Chalmers KJ, Smith JSC, Kilian A, et al. The genetic diversity of UK, US and Australian cultivars of Triticum aestivum measured by DArT markers and considered by genome. Theor Appl Genet. 2008;116:439–53.

21. Gardiner LJ, Brabbs T, Akhunov A, Jordan K, Budak H, Richmond T, et al. Integrating genomic resources to present full gene and putative promoter capture probe sets for bread wheat. Gigascience. Oxford University Press; 2019;8:1–13.

22. IWGSC. A chromosome-based draft sequence of the hexaploid bread wheat (Triticum aestivum) genome Ancient hybridizations among the ancestral genomes of bread wheat Genome interplay in the grain transcriptome of hexaploid bread wheat Structural and functional pa. Science [Internet]. 2014;345:1250092. Available from: http://www.sciencemag.org/content/345/6194/1250092.abstract

23. Kover PXPX, Valdar W, Trakalo J, Scarcelli N, Ehrenreich IMIMM, Purugganan MDMDD, et al. A Multiparent Advanced Generation Inter-Cross to fine-map quantitative traits in Arabidopsis thaliana. Plos Genet. 2009/07/14. 2009;5:e1000551.

24. Gan X, Stegle O, Behr J, Steffen JGG, Drewe P, Hildebrand KLL, et al. Multiple reference genomes and transcriptomes for Arabidopsis thaliana. Nature. 2011/08/30. 2011;477:419–23.

25. Camacho C, Coulouris G, Avagyan V, Ma N, Papadopoulos J, Bealer K, et al. BLAST+: Architecture and applications. BMC Bioinformatics. 2009;10:1–9.

26. Davies RW, Flint J, Myers S, Mott R. Rapid genotype imputation from sequence without reference panels. Nat Genet. 2016;48:965–9.

27. IWGSC. Shifting the limits in wheat research and breeding using a fully annotated reference genome. Science. 2018;361:eaar7191.

28. Cavanagh CR, Chao S, Wang S, Huang BE, Stephen S, Kiani S, et al. Genome-wide comparative diversity uncovers multiple targets of selection for improvement in hexaploid wheat landraces and cultivars. Proc Natl Acad Sci U S A. Proc Natl Acad Sci U S A; 2013;110:8057–62.

29. Broman KW. The genomes of recombinant inbred lines. Genetics. 2005;169:1133–46.

30. Pont C, Leroy T, Seidel M, Tondelli A, Duchemin W, Armisen D, et al. Tracing the ancestry of modern bread wheats. Nat Genet [Internet]. Springer US; 2019;51:905–11. Available from: http://dx.doi.org/10.1038/s41588-019-0393-z

31. Rhoné B, Raquin AL, Goldringer I. Strong linkage disequilibrium near the selected Yr17 resistance gene in a wheat experimental population. Theor Appl Genet. 2007;114:787–802.

32. Martynov SP, Dobrotvorskaya T V., Krupnov VA. Analysis of the Distribution of Triticum timopheevii Zhuk. Genetic Material in Common Wheat Varieties (Triticum aestivum L.). Russ J Genet. 2018;54:166–75.

33. Villareal RL, Toro E, Mujeeb-Kazi A, Rajaram S. The 1BL/1RS chromosome translocation effect on yield characteristics in a Triticum aestivum L. cross. Plant Breed. John Wiley & Sons, Ltd; 1995;114:497–500.

34. Tsilo TJ, Jin Y, Anderson JA. Diagnostic Microsatellite Markers for the Detection of Stem Rust Resistance Gene *Sr36* in Diverse Genetic Backgrounds of Wheat. Crop Sci. John Wiley & Sons, Ltd; 2008;48:253–61.

35. Gardner KA, Wittern LM, Mackay IJ. A highly recombined, high-density, eight-founder wheat MAGIC map reveals extensive segregation distortion and genomic locations of introgression segments. Plant Biotechnol J. 2016;14:1406–17.

36. Robert O, Abelard C, Dedryver F. Identification of molecular markers for the detection of the yellow rust resistance gene Yr17 in wheat. Mol Breed. 1999;5:167–75.

37. Marchal C, Zhang J, Zhang P, Fenwick P, Steuernagel B, Adamski NM, et al. BED-domain-containing immune receptors confer diverse resistance spectra to yellow rust. Nat Plants [Internet]. Springer US; 2018;4:662–8. Available from: http://dx.doi.org/10.1038/s41477-018-0236-4

38. Xu D, Wen W, Fu L, Li F, Li J, Xie L, et al. Genetic dissection of a major QTL for kernel weight spanning the Rht-B1 locus in bread wheat. Theor Appl Genet [Internet]. Springer Berlin Heidelberg; 2019;132:3191–200. Available from: https://doi.org/10.1007/s00122-019-03418-w

39. Zhang M, Gao M, Zheng H, Yuan Y, Zhou X, Guo Y, et al. QTL mapping for nitrogen use efficiency and agronomic traits at the seedling and maturity stages in wheat. Mol Breed. Molecular Breeding; 2019;39.

40. Kuzay S, Xu Y, Zhang J, Katz A, Pearce S, Su Z, et al. Identification of a candidate gene for a QTL for spikelet number per spike on wheat chromosome arm 7AL by high-resolution genetic mapping. Theor Appl Genet [Internet]. Springer Berlin Heidelberg; 2019;132:2689–705. Available from: https://doi.org/10.1007/s00122-019-03382-5

41. Wang D, Yu K, Jin D, Sun L, Chu J, Wu W, et al. Natural variations in the promoter of Awn Length Inhibitor 1 (ALI-1) are associated with awn elongation and grain length in common wheat. Plant J. 2020;101:1075–90.

42. Lin M, Zhang D, Liu S, Zhang G, Yu J, Fritz AK, et al. Genome-wide association analysis on pre-harvest sprouting resistance and grain color in U.S. winter wheat. BMC Genomics [Internet]. BMC Genomics; 2016;17. Available from: http://dx.doi.org/10.1186/s12864-016-3148-6

43. Beales J, Turner A, Griffiths S, Snape JW, Laurie DA. A pseudo-response regulator is misexpressed in the photoperiod insensitive Ppd-D1a mutant of wheat (Triticum aestivum L.). Theor Appl Genet. Theor Appl Genet; 2007;115:721–33.

44. Kamran A, Iqbal M, Spaner D. Flowering time in wheat (Triticum aestivum L.): A key factor for global adaptability. Euphytica. 2014;197:1–26.

45. Hedden P. The genes of the Green Revolution. Trends Genet. 2003;19:5–9.

46. de Los Campos G, Hickey JM, Pong-Wong R, Daetwyler HD, Calus MPL. Whole-genome regression and prediction methods applied to plant and animal breeding. Genetics. Genetics; 2013;193:327–45.

47. Michel S, Löschenberger F, Ametz C, Pachler B, Sparry E, Bürstmayr H. Combining grain yield, protein content and protein quality by multi-trait genomic selection in bread wheat. Theor Appl Genet [Internet]. Springer Berlin Heidelberg; 2019;132:2767–80. Available from: https://doi.org/10.1007/s00122-019-03386-1

48. Michel S, Löschenberger F, Ametz C, Pachler B, Sparry E, Bürstmayr H. Simultaneous selection for grain yield and protein content in genomics-assisted wheat breeding. Theor Appl Genet [Internet]. Springer Berlin Heidelberg; 2019;132:1745–60. Available from: https://doi.org/10.1007/s00122-019-03312-5

49. Gan X, Stegle O, Behr J, Steffen JGJGG, Drewe P, Hildebrand KLLKL, et al. Multiple reference genomes and transcriptomes for Arabidopsis thaliana. Nature. 2011/08/30. 2011;477:419–23.

50. Borrill P, Harrington SA, Uauy C. Applying the latest advances in genomics and phenomics for trait discovery in polyploid wheat. Plant J. 2019;97:56–72.

51. Fradgley N, Gardner KA, Cockram J, Elderfield J, Hickey JM, Howell P, et al. A large-scale pedigree resource of wheat reveals evidence for adaptation and selection by breeders. PLoS Biol. 2019;17:1–20.

52. van de Wouw M, van Hintum T, Kik C, van Treuren R, Visser B. Genetic diversity trends in twentieth century crop cultivars: A meta analysis. Theor Appl Genet. 2010;120:1241–52.

53. Fu YB. Understanding crop genetic diversity under modern plant breeding. Theor Appl Genet. Springer Berlin Heidelberg; 2015;128:2131–42.

54. Baud A, Hermsen R, Guryev V, Stridh P, Graham D, McBride MWMW, et al. Combined sequence-based and genetic mapping analysis of complex traits in outbred rats. Nat Genet. 2013;45:767–75.

55. Bogard M, Allard V, Brancourt-Hulmel M, Heumez E, Machet J-M, Jeuffroy M-H, et al. Deviation from the grain protein concentration–grain yield negative relationship is highly correlated to post-anthesis N uptake in winter wheat. J Exp Bot. Oxford Academic; 2010;61:4303–12.

56. Cormier F, Faure S, Dubreuil P, Heumez E, Beauchêne K, Lafarge S, et al. A multi-environmental study of recent breeding progress on nitrogen use efficiency in wheat (Triticum aestivum L.). Theor Appl Genet. 2013;126:3035–48.

57. Lenz PRN, Beaulieu J, Mansfield SD, Clément S, Desponts M, Bousquet J. Factors affecting the accuracy of genomic selection for growth and wood quality traits in an advanced-breeding population of black spruce (Picea mariana). BMC Genomics. BMC Genomics; 2017;18:1–17.

58. Moser G, Khatkar MS, Hayes BJ, Raadsma HW. Accuracy of direct genomic values in Holstein bulls and cows using subsets of SNP markers. Genet Sel Evol. 2010;42:1–15.

59. Duncan L, Shen H, Gelaye B, Meijsen J, Ressler K, Feldman M, et al. Analysis of polygenic risk score usage and performance in diverse human populations. Nat Commun. Nature Publishing Group; 2019;10:3328.

60. Liu K, Muse S V. PowerMaker: An integrated analysis environment for genetic maker analysis. Bioinformatics. 2005;21:2128–9.

61. Porras-Hurtado L, Ruiz Y, Santos C, Phillips C, Carracedo Á, Lareu M V. An overview of STRUCTURE: Applications, parameter settings, and supporting software. Front Genet. 2013;4:1–13.

62. Mackay IJ, Bansept-Basler P, Bentley AR, Cockram J, Gosman N, Greenland AJ, et al. An eight-parent multiparent advanced generation inter-cross population for winter-sown wheat: Creation, properties, and validation. G3 Genes, Genomes, Genet. 2014;4:1603–10.

63. Butler D, Cullis B, Gilmour A, Gogel B. ASReml–R Reference Manual [Internet]. Brisbane: The State of Queensland, Department of Primary Industries and Fisheries; 2009. Available from: http://scholar.google.com/scholar?hl=en&btnG=Search&q=intitle:mixed+models+for+S+language+environments+ASReml-R+reference+manual#0

64. Li H, Durbin R. Fast and accurate short read alignment with Burrows-Wheeler transform. Bioinformatics. 2009/05/20. 2009;25:1754–60.

65. Li H, Handsaker B, Wysoker A, Fennell T, Ruan J, Homer N, et al. The Sequence Alignment/Map format and SAMtools. Bioinformatics. 2009/06/10. 2009;25:2078–9.

66. McKenna A, Hanna M, Banks E, Sivachenko A, Cibulskis K, Kernytsky A, et al. The Genome Analysis Toolkit: a MapReduce framework for analyzing next-generation DNA sequencing data. Genome Res. 2010;20:1297–303.

67. Li H. A statistical framework for SNP calling, mutation discovery, association mapping and population genetical parameter estimation from sequencing data. Bioinformatics. 2011;27:2987–93.

68. R Core Team. R: A Language and Environment for Statistical Computing. Vienna, Austria; 2016.

69. Kang HM, Zaitlen NA, Wade CM, Kirby A, Heckerman D, Daly MJ, et al. Efficient control of population structure in model organism association mapping. Genetics. 2008/04/04. 2008;178:1709–23.

70. VanRaden P. Efficient estimation of breeding values from dense genomic data. J Dairy Sci. 2007;90:374–5.

71. Habier D, Fernando RL, Dekkers JCM. The impact of genetic relationship information on genome-assisted breeding values. Genetics. 2007;177:2389–97.

72. Friedman J, Hastie T, Tibshirani R. glmnet: Lasso and elastic-net regularized generalized linear models. R Packag version 14. 2009;

