## Supplementary Figures for "Limited haplotype diversity underlies polygenic trait architecture across 70 years of wheat breeding"

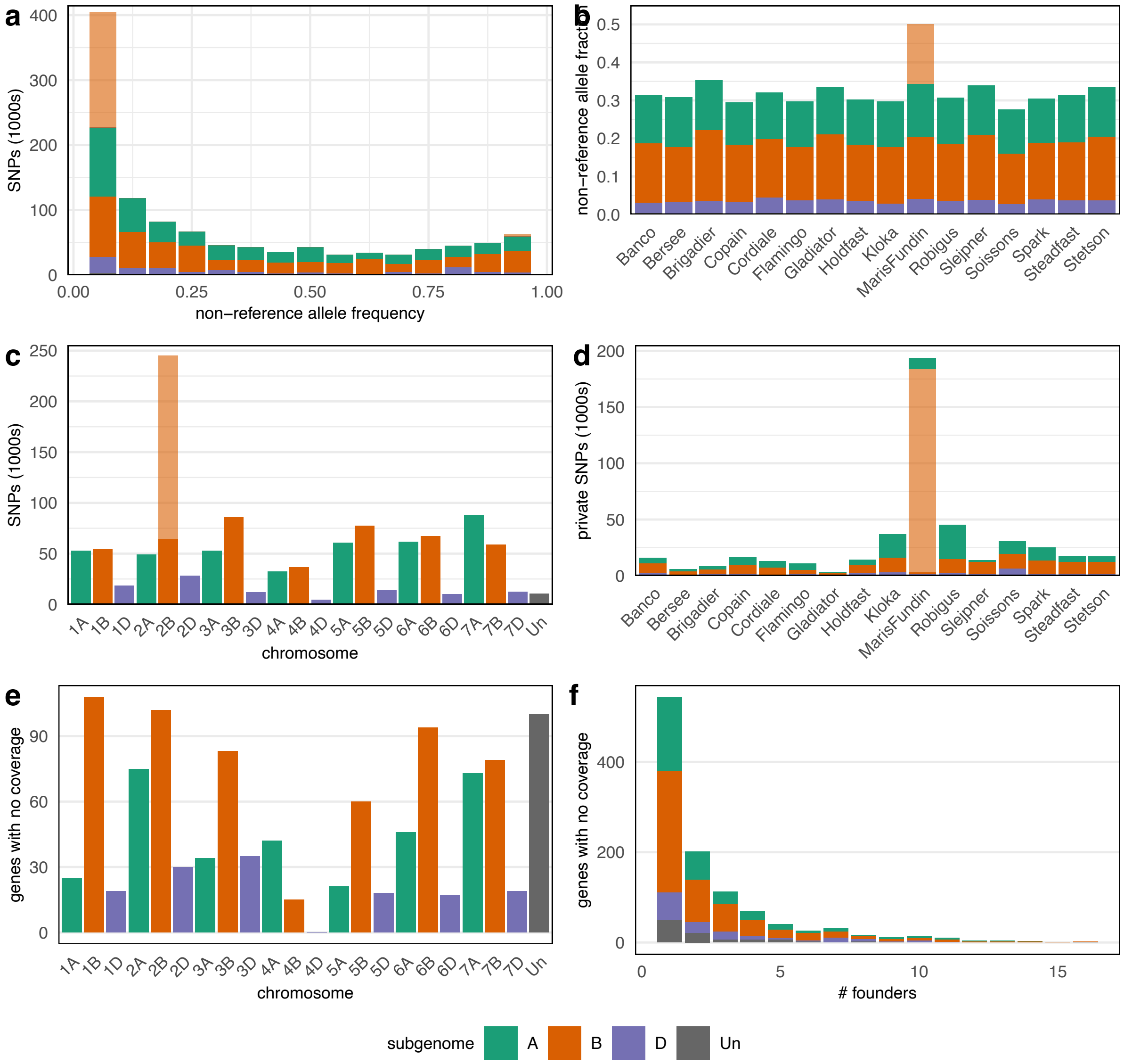


*Supplementary Figure 1: Distribution of SNPs across chromosomes and founders. (a) shows the non reference allele frequency spectrum, (b) shows the proportion of non-reference alleles called in each founder, (c) shows the chromosomal distribution of SNPs and (d) shows the distribution of private SNPs (minor allele found in only one founder) across founders. There are a large number of SNPs that are private to Maris Fundin and found on chromosome 2B. There are likely due to an introgression from T. timopheevi and are indicated using a paler shade throughout. (e) shows the fraction of gene-promoter pairs that have no coverage in at least one founder by chromosome. (f) shows the number of founders that are missing alignments for particular gene-promoter pairs. Colours indicate subgenome locations, with fewer SNPs segregating on the D subgenome, which was acquired in the most recent allo-polyploidisation event.*

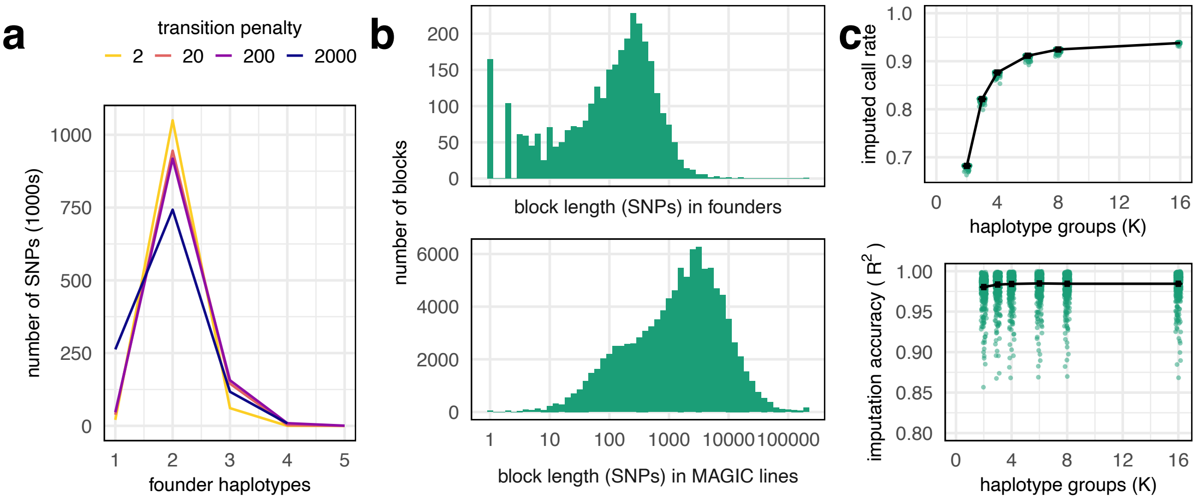


*Supplementary Figure 2: Haplotype block length in founders and MAGIC lines. Founder haplotype blocks were determined using a dynamic programming algorithm that compares the similarity between all pairs of founders. (a) For various transition penalty parameters, two haplotypes are inferred across most of the genome. In the main text, we refer to the transition penalty 200. (b) The block lengths (in SNP numbers) in the founders are smaller than those of the MAGIC lines. In MAGIC lines, haplotype blocks are defined as consecutive SNPs that are inferred to have the same founder ancestry (>90% dosage). (c) Imputation call rate and accuracy when assuming that MAGIC line mosaics are defined by a different numbers of haplotype groups.*

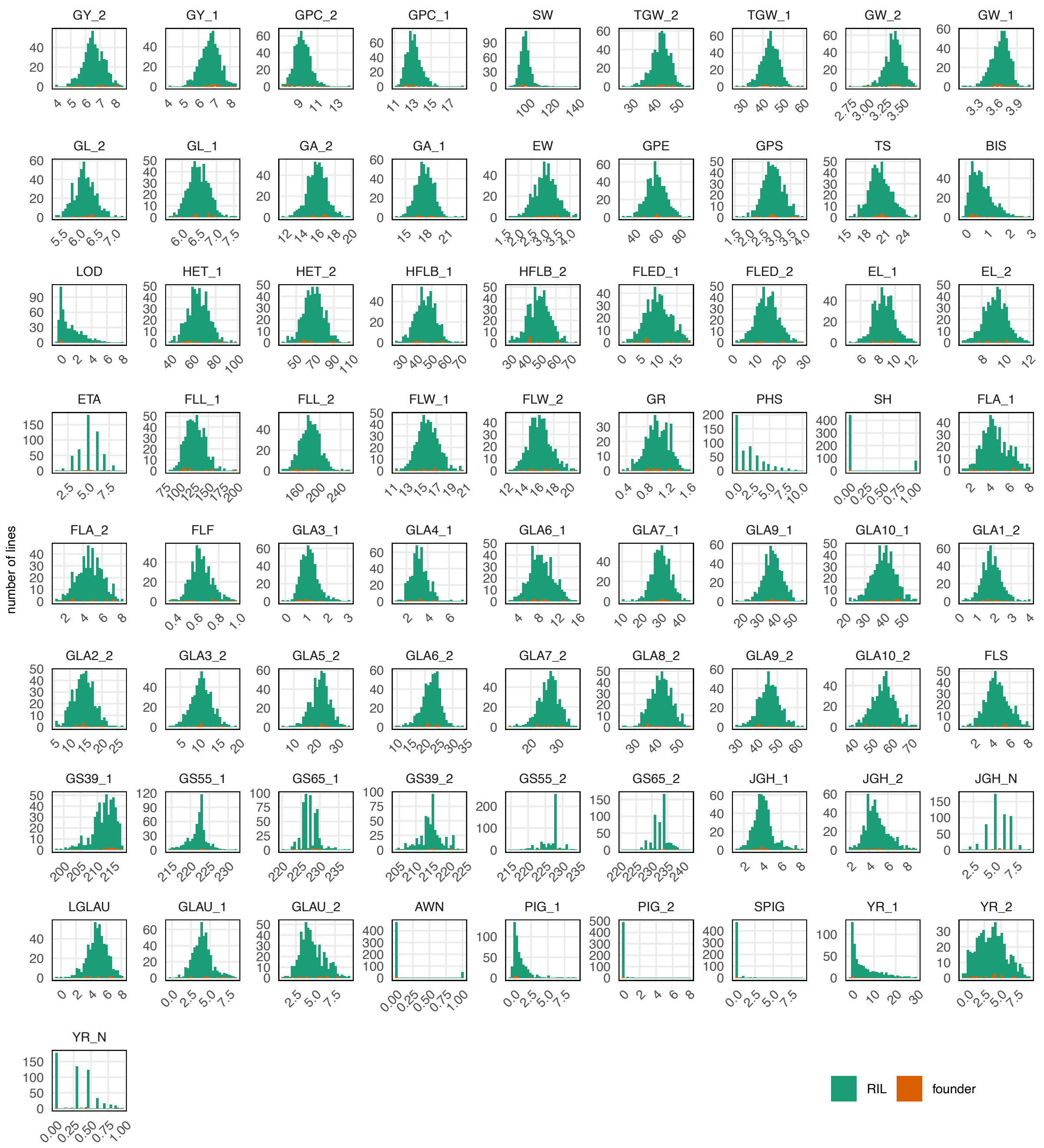


*Supplementary Figure 3 Distribution of Best Linear Unbiased Estimates (BLUEs) for founders (orange) and the RILs derived from them (green) across 73 phenotypic measurements. For most normally-distributed phenotypes, the phenotypic range of the RILs is wider than that of the founders (transgressive segregation).*

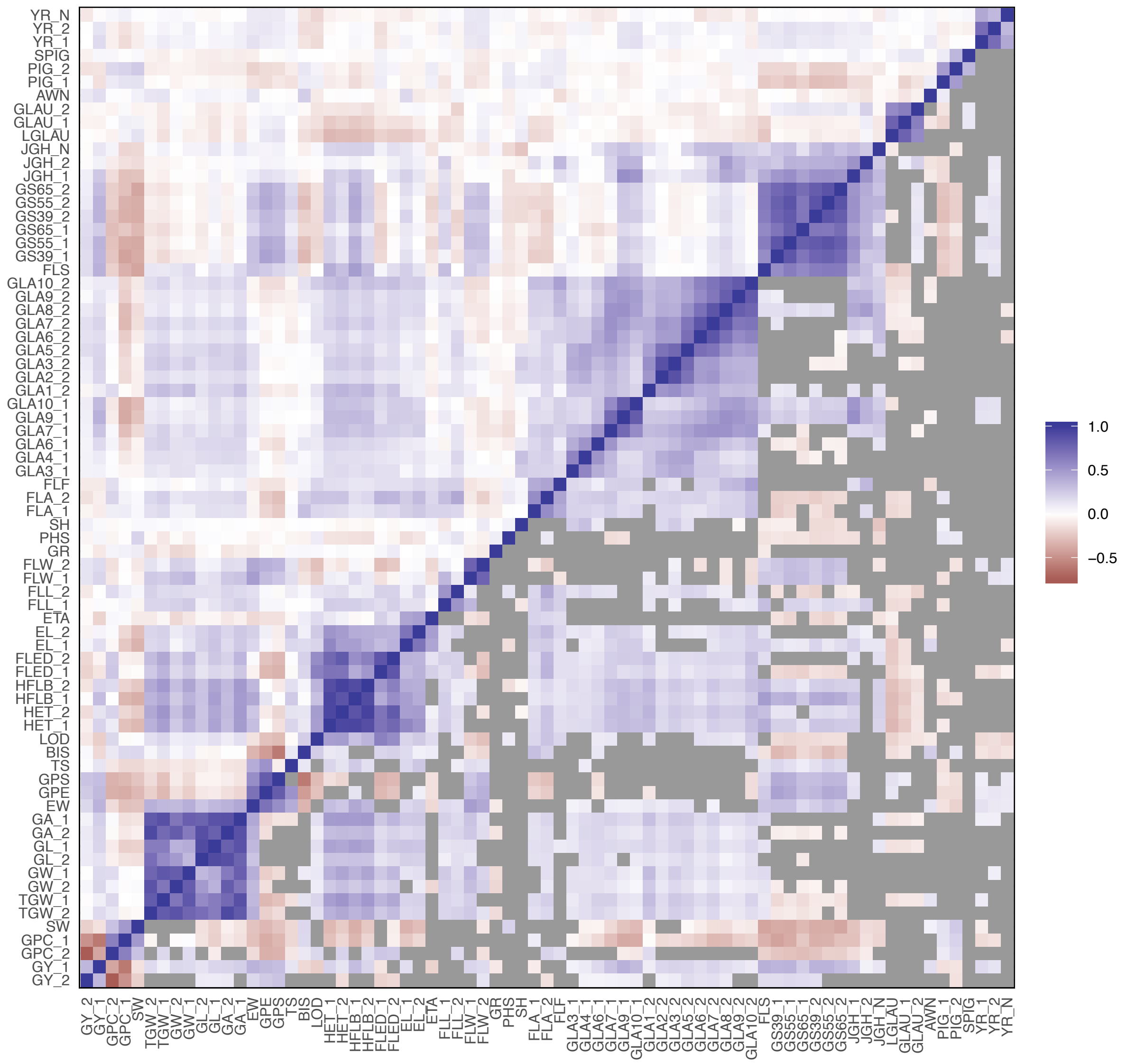

*Supplementary Figure 4 Correlations between phenotypes in the 504 RILs. The heatmap fill colour shows the Pearson's correlation coefficient for all pairwise combinations of phenotype measurements. In the lower diagonal, all non-significant correlations (p>0.05 for the null hypothesis of zero correlation) are grey.*

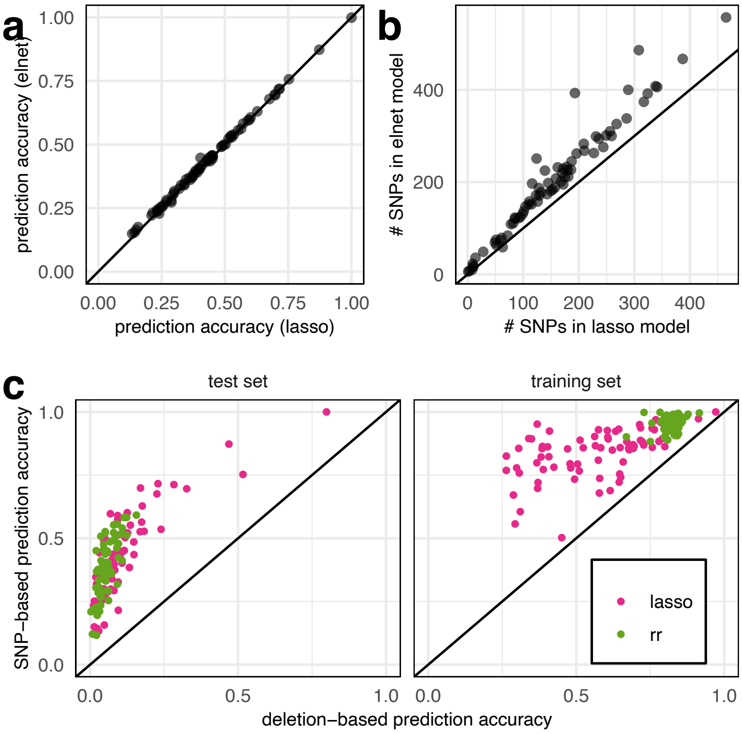


*Figure 5 Comparison between genomic prediction using LASSO and ELNET (a,b) and between genomic prediction based on SNPs and gene deletion scores (c). SNP-based prediction accuracy using LASSO and ELNET are almost identical (a), based on the mean correlation coefficient between predictions and phenotypes for the test set of lines across 50 cross-validation replicates. However, the LASSO prediction models include far fewer SNPs (b), where we show the number of SNPs included in the full model trained on all 504 RILs. (c) shows that the prediction accuracy of models using gene deletion scores never exceeds that of SNP-based prediction.*
